# Social Factors of Health Covary with Population Stratification and Confound Heritability Estimates

**DOI:** 10.1101/2025.06.12.659317

**Authors:** Jaclyn L. Liquori, Oona Shigeno Risse-Adams, Rangarajan Bharadwaj, Dylan Ong, Nasa Sinnott-Armstrong, Shaila A. Musharoff

**Author notes:** These authors contributed equally to this work (Jaclyn L. Liquori, Oona Shigeno Risse-Adams, Nasa Sinnott-Armstrong, Shaila A. Musharoff).

## Abstract

Social factors of health (SFOH) are critical determinants of human traits but are rarely incorporated into genetic models. Here, we assess how participant-completed SFOH survey data influence heritability estimates and trait variance in 85,963 diverse All of Us participants spanning multiple ancestries and racial identities. We summarized SFOH survey data using multiple correspondence analysis (MCA) into axes representing distinct social dimensions (e.g., social support, perceived stress) and developed a Social Similarity Index (SSI) to capture social environment similarity among individuals. To analyze trait heritability, we used Haseman-Elston (HE) regression for its sensitivity to residual structure. Including SFOH measures as covariates in HE regression models significantly reduced heritability estimates for four of 18 traits, three of which are anthropometric: body mass index, hip circumference, waist circumference, and HDL cholesterol. This suggests that unmodeled social structure can be misattributed to genetic effects. Though several SFOH measures have non-zero heritability when adjusting for three genetic principal components (*h*^2^ *≈* 0.01*−*0.09), these are reduced to similar estimates when adjusting for seven genetic principal components (*h*^2^ *≈* 0.01*−*0.02). This convergence across SFOH measures, which capture different social dimensions, indicates that SFOH measures capture population stratification, emphasizing their utility for reducing confounding in heritability estimates. SFOH measures were also associated with trait variance, with anthropometric traits—those with reduced heritability—exhibiting the most extensive dispersion effects. This implies that social environment influences trait variability. Our findings highlight the necessity of including social and environmental factors in genetic studies to reduce potential confounding.

## 1 Introduction

Modeling the social drivers of human traits remains a critical challenge in biomedicine. This is particularly acute in statistical genetics, where genome-wide studies routinely adjust for genetic stratification [1–4], while trait-relevant social and environmental exposures typically remain unmeasured or oversimplified. Social factors of health (SFOH)—such as housing, financial insecurity, education, and perceived stress—shape both disease risk and access to health care [5–9]. SFOH are increasingly captured in structured survey form in biobank-scale cohorts, including the All of Us Research Program [10]. However, SFOH data are rarely explicitly incorporated in heritability and genetic association models [11, 12]. Like genetic variation, these factors are stratified within populations but can change rapidly and profoundly impact health outcomes. This omission can leave residual trait variation structured by social factors, such that social stratification is misattributed as genetic variance, creating incomplete inferences about underlying genetic architecture shaped by spurious signals. Although the All of Us Social Determinants of Health (SDOH) survey data has been previously characterized [13] and connected to health outcomes [14, 15], to our knowledge, it has not been incorporated in heritability models.

Survey data often contain trait-relevant information, but their relative structure and information content differ from those of genetic [16] data and from the assumptions of standard genetic models, making joint modeling challenging. A recent work applied PCA to dietary survey data and used PC1 as an environmental factor in genetic association testing [17]. Dual likelihood modeling frameworks [18] can address some of these technological challenges. Furthermore, survey data have known participation bias [19] as well as completion and non-response rates that are correlated with demographic factors, such as race and ethnicity [20]. Because survey completion rates vary by race and ethnicity, it becomes difficult to determine whether observed patterns reflect true population-specific factors or simply artifacts of differential survey participation. This challenge is particularly acute in multi-ancestry studies. Related methodological work has addressed the need for ancestry-aware approaches when modeling complex traits [21].

Stratifying analyses by self-identified race or ethnicity can itself be problematic, as these social constructs overlap substantially with genetic ancestry patterns yet do not map cleanly onto them, creating challenges in accurately attributing variance components [22–24]. Recent methodological advances have developed extensions of Haseman-Elston regression for estimating heritability to account for population stratification while assuming constant heritability across subpopulations [25]. However, this assumption might not hold when social and genetic stratification are entangled, as we investigate here. Rather than attempting to disentangle these overlapping signals through stratification, we conduct our analysis in the multi-ancestry and multi-racial group of 85,963 participants, allowing us to characterize how SFOH and genetic factors jointly structure trait variation across the social and genetic diversity of these individuals. SFOH survey data contains information that is potentially correlated with genetic data, and understanding the relationship between the two is critical for robust joint analysis while avoiding additional bias in genetic estimates.

Here, we investigate how high-dimensional social structure captured by the All of Us SDOH survey influences single nucleotide polymorphism-based heritability (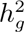)—the proportion of phenotypic variance in a population explained by additive genetic effects of common SNPs—of 18 anthropometric and metabolic traits. We use the terms “heritability” and *h*^2^ to refer to this SNP-based heritability (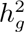), which provides an estimate for narrow-sense heritability. In this paper, we address three specific questions: (1) Do SFOH measures themselves have non-zero heritability, suggesting potential confounding between social and genetic structures? (2) Does accounting for SFOH reduce trait heritability estimates, indicating that social structure may be misattributed to genetic effects? (3) Do SFOH measures influence trait variance patterns beyond their effects on means? To address these questions, we summarize the SDOH survey responses into SFOH measures using two complementary strategies: dimensionality reduction via multiple correspondence analysis (MCA) and a Social Similarity Index (SSI) derived from an environmental relatedness matrix (ERM). These yield participant-level covariates that integrate the critical yet often overlooked influences of social environment factors and facilitate their joint analysis with genetic factors.

## 2 Results

We analyzed data from 85,963 All of Us (v7) participants who met the following criteria: availability of SDOH survey data, genotyping array data, a selected sex at birth of “female” or “male” (Table S1), and a self-identified race (Table S2) from The Basics survey. We performed analyses of collinearity and the relationship between social and population genetic structure on this combined analysis group, while heritability estimation was restricted to subsets of individuals with measurements for each of the 18 analyzed quantitative traits.

### 2.1 Collinearity

To characterize the relationship between social and genetic measures, we first derived two sets of SFOH measures from one-hot encoded All of Us SDOH survey data: the SFOH SSI and SFOH MC axes. We then assessed collinearity by calculating Spearman correlations between these SFOH MC axes and genetic principal components (PCs) (Fig. 1). For the SSI, the maximum absolute Pearson correlation with any genetic PC was 0.317 and all variance inflation factors (VIFs), where values above 5 signal moderate multicollinearity, remained below 1.150, indicating an acceptable amount of multicollinearity. For the top forty SFOH MC axes, the maximum correlation with any genetic PC was 0.011 (VIFs *<* 1.145). Thus, each SFOH measure can safely be modeled alongside genetic PCs without problematic multicollinearity.

**Fig. 1:**
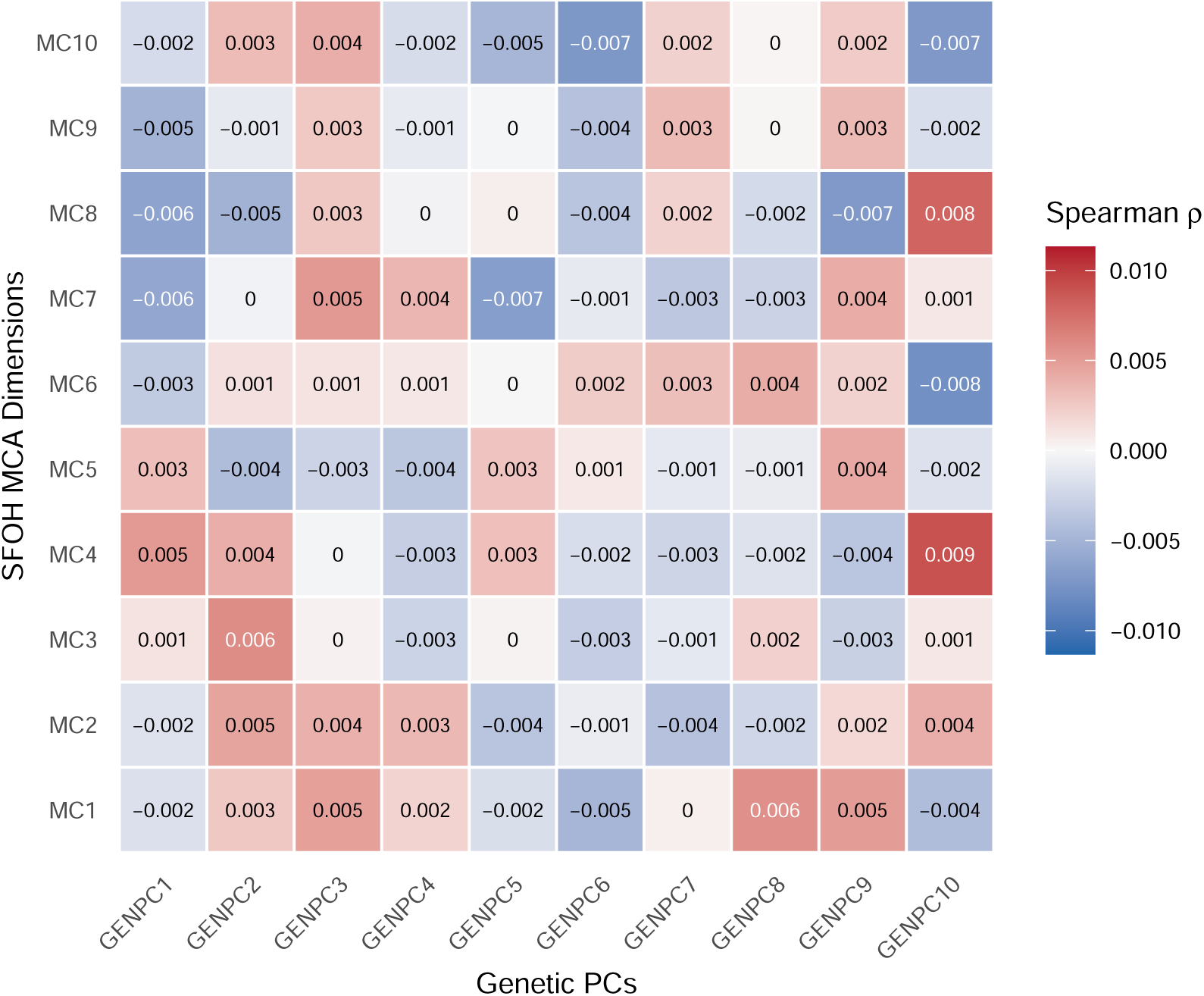
Relationship between genetic structure and SFOH measures. Spearman correlations between the 10 genetic PCs (x-axis) and top 10 SFOH survey derived MC axes (y-axis).

### 2.2 Missingness Analysis

We excluded missing responses coded as “PMI: Skip”, “PMI: Don’t Know”, “PMI: Not Sure”, “PMI: Other” or “PMI: Prefer Not To Answer”. At the question-level, missingness was very low (less than 5%) for nearly all themed items (Table S3), with only two un-themed questions exhibiting high skip rates. Theme-level missingness was similar across self-identified race groups (2–8%) (Table S4), and at the individual level the median per-participant missingness was approximately 3%, with less than 1% of participants missing more than 20% of items.

### 2.3 Trait heritability

Using GCTA [26], we estimate heritability using Haseman–Elston (HE) regression [27], a computationally efficient method of moments estimator suitable for large datasets. We chose HE regression over the genomic-relatedness-based restricted maximum-likelihood (GREML) approach because its sensitivity to residual structure makes it well-suited for detecting when SFOH measures capture variance that would otherwise be misattributed to genetic effects—our primary research question. To assess how SFOH measures and genetic PCs influence these estimates, we tested six nested covariate models (Table 1) that progress from basic demographic adjustment—age, sex, self-identified race, and self-identified ethnicity (model 1)—to full covariate sets (model 6). All analyses of systolic blood pressure include adjustment for medication use. The single-statistic SFOH SSI only appears in two models (2, 5) while the partitioned multi-axis SFOH MCA appears in four models (2, 3, 5, and 6). The lead SFOH MC axes 1:6 (14.73% variance explained) (Fig. S1), appear in models 2 and 5, while the comprehensive SFOH MC axes 1:40 (38.02% variance explained), appear in models 3 and 6. The top genetic PCs 1:7 (96.84% of variance explained) (Fig. S2) are used in models 4, 5, and 6.

**Table 1:** Covariate models used in trait heritability analyses. Models build progressively from no SFOH adjustment to full SFOH adjustment.

| SFOH Adjustment Model | SFOH Covariates |
| --- | --- |
| 1. No Adjustment | None |
| 2. Lead Measures | SFOH MC axes 1:6 or SFOH SSI |
| 3. Comprehensive Measures | SFOH MC axes 1:40 |
| 4. No Adjustment + Genetic PCs 1:7 | None |
| 5. Lead Measures + Genetic PCs 1:7 | SFOH MC axes 1:6, or SFOH SSI |
| 6. Comprehensive Measures + Genetic PCs 1:7 | SFOH MC axes 1:40 |

#### 2.3.1 HE regression is sensitive to unmodeled SFOH

We used HE regression to estimate heritability via cross product (HE-CP). In the combined analysis group of up to 85,963 individuals (Fig. 2), adding genetic PCs 1:7 (models 4, 5, and 6) consistently reduced heritability estimates relative to the Base (model 1) and SFOH-only models (models 2 and 3). These reductions were significant in 11 of 18 traits: BMI, HDL cholesterol, carbon dioxide, creatinine, diastolic blood pressure, glucose, hip circumference, heart rate, sodium, urea nitrogen, and waist circumference. Sex-stratified analyses revealed similar patterns, with minor variation in estimates between females (Fig. S3) and males (Fig. S4) for certain traits. This pattern indicates that genetic PCs capture population structure that inflates heritability estimates when unaccounted for, which is a known property of HE regression. When genetic ancestry is associated with trait values, this method may misattribute this covariance to genetic factors; including genetic PCs as covariates corrects for this potential source of confounding.

**Fig. 2:**
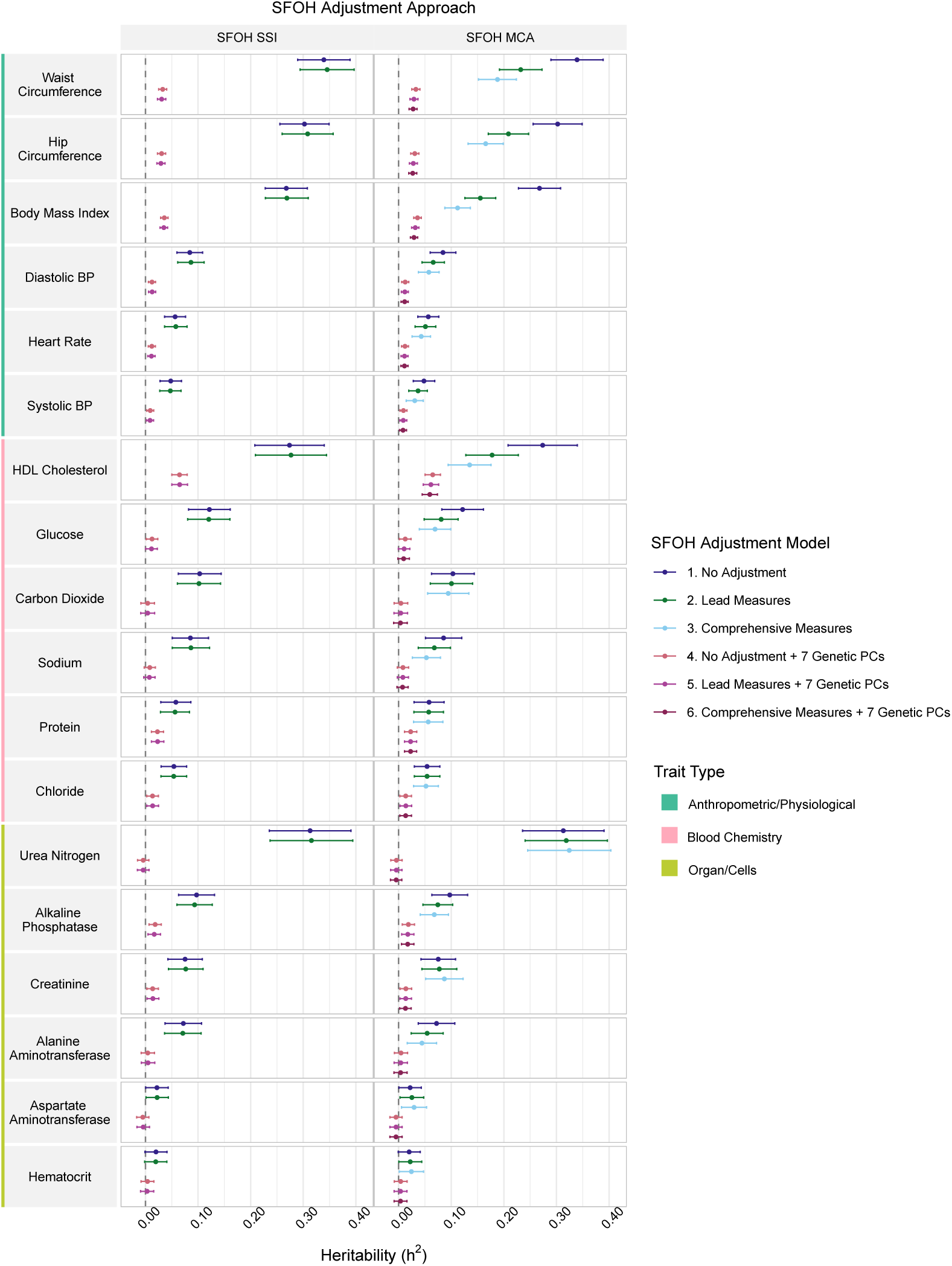
Combined analysis group Haseman-Elston (HE) regression across SFOH-covariate models. HE regression heritability estimates for all relevant participants with 95% confidence intervals for 18 traits (rows), calculated using two Social Factors of Health (SFOH) summarization strategies (columns). All models are adjusted for age, sex, race, and ethnicity. Models with SFOH adjustment include lead (MCs 1:6 or SSI) or comprehensive (MCs 1:40) adjustment. Adding SFOH MC axes (models 2,3) reduces heritability estimates for body mass index, HDL cholesterol, hip circumference, glucose, and waist circumference relative to the base model (model 1). Points represent HE-CP estimates; intervals are 95% CIs from jackknife standard errors. The dashed line indicates a heritability of zero.

Adding SFOH MC axes to the Base model (models 2 and 3) also reduced the heritability estimate of 5 of 18 traits-BMI, hip circumference, waist circumference, glucose, and HDL cholesterol (Fig. 3)—while the SSI had no significant effect on any trait (Table S5). This suggests that a global measure of social similarity is less consequential for heritability estimation than specific social dimensions. Specifically, including SFOH MC axes 1:6 (model 2) significantly reduced heritability estimates relative to the Base model (model 1) for BMI (by 0.113 from 0.268 to 0.155), hip circumference (by 0.093 from 0.302 to 0.209), and waist circumference (by 0.107 from 0.339 to 0.232). Including the MC axes 7:40 (model 3) further reduced the heritability of these traits, though not significantly (Table S6). As a sensitivity analysis, we also performed PCA on the survey data [17], here we use the same one-hot encoded SFOH survey data. When including SFOH PCs into heritability models, results were broadly consistent with MCA findings (Fig. S5).

**Fig. 3:**
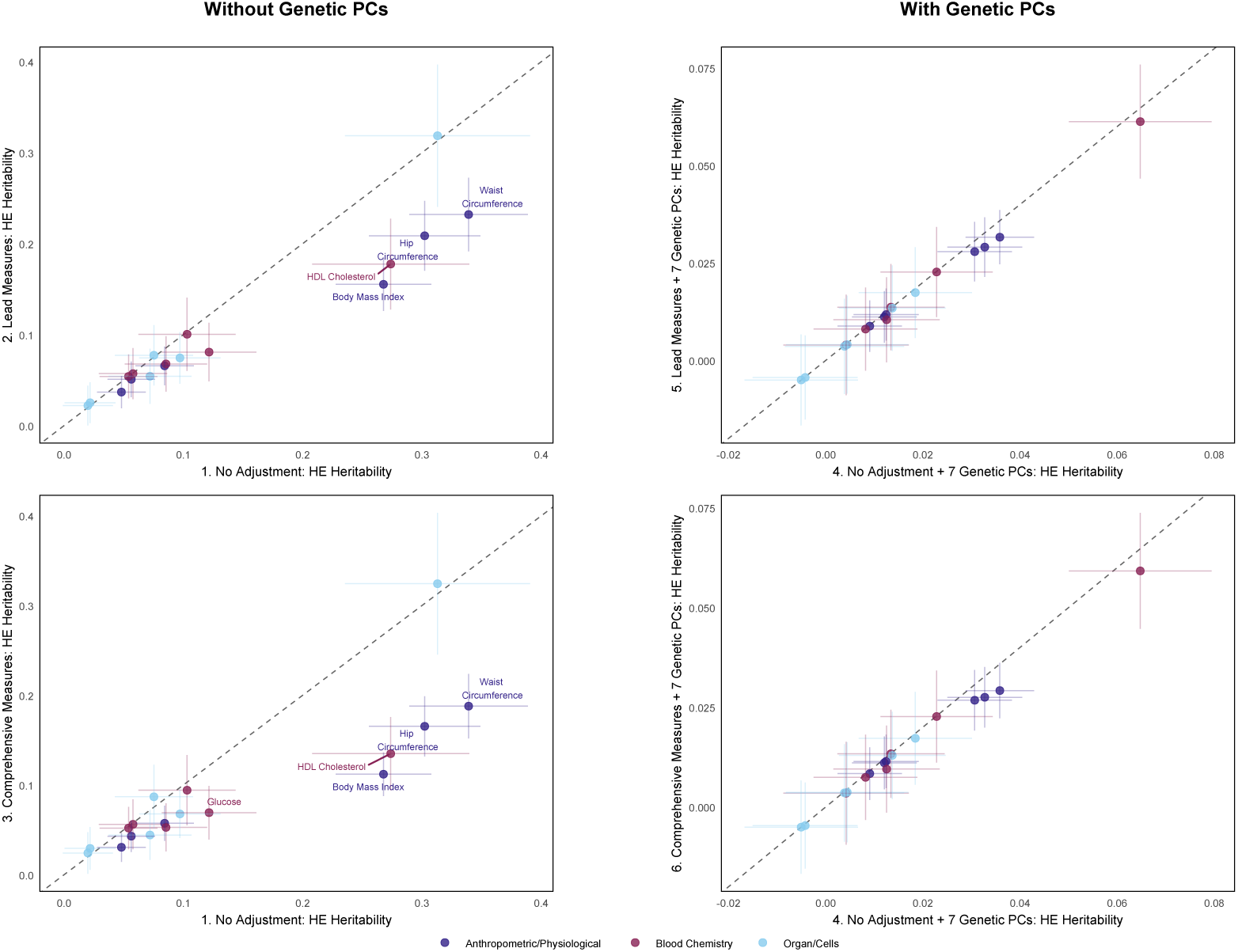
Combined analysis group Haseman-Elston (HE) regression across SFOH MCA adjustment models. HE regression heritability estimates for all relevant participants with 95% confidence intervals for 18 traits, calculated using SFOH MCA. Panels are arranged in a 2×2 grid: models without genetic PCs (left) and with genetic PCs (right). Each panel shows a pairwise model comparison with model numbers noted at the beginning of the x- and y-axis labels. All models are adjusted for age, sex, race, and ethnicity. Models with SFOH adjustment include lead (MCs 1:6) or comprehensive (MCs 1:40) adjustment. Adding SFOH MC axes (models 2,3) reduces heritability estimates for BMI, HDL cholesterol, hip circumference, glucose, and waist circumference relative to the base model (model 1). Points represent HE-CP estimates; intervals are 95% CIs from jackknife standard errors. The dashed line indicates the y = x line.

The sensitivity of HE regression to SFOH adjustment is consistent with SFOH measures capturing social structure that correlates with genetic similarity. Our finding that specific SFOH measures initially appear heritable but decrease with comprehensive genetic PC adjustment (Section 2.4) indicates this correlation primarily reflects population stratification rather than true genetic effects on social factors. Because HE regression attributes any phenotypic covariance aligned with the GRM to additive genetic variance, SFOH measures that proxy genetic stratification can substantially influence estimates when included as covariates. Including SFOH measures removes this shared variance, leading to lower *h*^2^ estimates even without genetic PCs. This demonstrates that SFOH measures can act as proxies for confounding population structure that would otherwise be misattributed to genetic effects.

These analyses demonstrate that survey-derived SFOH data summarized with MCA can influence heritability estimates obtained via HE regression, sometimes significantly so. Our results suggest that genetic PCs and SFOH measures capture overlapping information about population stratification, with SFOH measures potentially providing additional adjustment for residual structure not fully captured by a limited number of genetic PCs. We observed a similar pattern in an ancestry-specific analysis restricted to 48,146 individuals of European ancestry based on their genetic similarity to European reference populations in gnomAD, HGDP, and 1000 Genomes who additionally identified as “White” and “Not Hispanic or Latino”, which we refer to as the “European-ancestry subgroup”. For this analysis, the SFOH MC axes were recalculated using only the survey data from this subgroup of individuals. Despite reduced statistical power and larger standard errors, the pattern seen in the combined group analysis persisted (Fig. S6), suggesting that the confounding effect of social structure is not an artifact of multi-ancestry analysis alone.

### 2.4 SFOH measures show progressive reduction in heritability with genetic PC adjustment

To assess whether the SFOH measures are estimated to have non-zero heritability, we applied HE regression the SFOH MC axes and the SSI as traits. We tested three nested covariate models (Table 2) to assess how basic demographics and genetic PCs influence these estimates.

**Table 2:** Covariate models used in SFOH measure heritability analyses.

| Model | Covariates |
| --- | --- |
| 1. | Base: Age + Sex + Race + Ethnicity |
| 2. | Base + Genetic PCs 1:3 |
| 3. | Base + Genetic PCs 1:7 |

#### 2.4.1 SFOH heritability estimates decrease with comprehensive genetic PC adjustment

Initial heritability estimates for many SFOH measures were significantly different from zero when adjusting only for basic demographics (Fig. S7 and Fig. S8), some substantially so with estimates exceeding *h*^2^ = 0.600. However, these estimates decreased dramatically upon inclusion of genetic PCs. For MC5, heritability decreased substantially (from *h*^2^ = 0.086 to 0.013) when increasing adjustment from three to seven genetic PCs. Notably, with seven PCs, heritability estimates for all major SFOH measures converged to a similarly low baseline (Fig. 4, Table S7). We observed a similar pattern in the European-ancestry subgroup using the subgroup-specific MC axes (Fig. S9), though with reduced statistical power, reinforcing that the non-zero heritability of SFOH MCs primarily reflects population stratification, which operates both between and within ancestral groups. The latent structure captured by MCA showed strong consistency between the multi-ancestry combined analysis group MC axes and the European-ancestry subgroup MC axes (Fig. S10), with corresponding dimensions exhibiting moderate to high correlations.

**Fig. 4:**
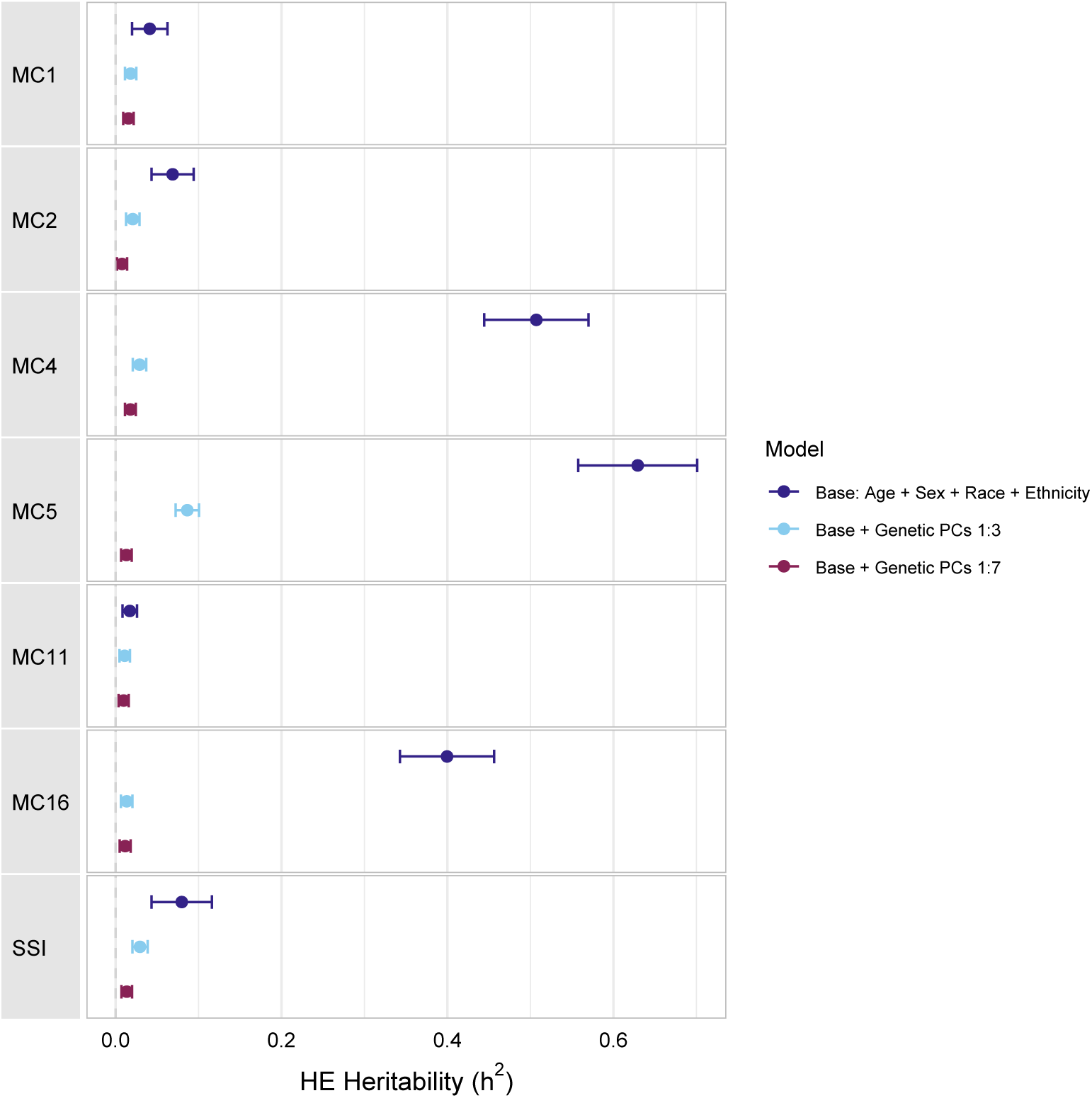
Combined analysis group Haseman-Elston (HE) regression heritability of SFOH measures that remain heritable after including seven genetic PCs. HE regression heritability estimates for all relevant participants with 95% confidence intervals for SFOH measures that appear heritable after comprehensive genetic PC adjustment. Points represent HE-CP estimates; intervals are 95% CIs from jackknife standard errors. The dashed line indicates a heritability of zero. HE regression heritability estimates show progressive reduction with genetic PC adjustment, demonstrating that apparent heritability primarily reflects population stratification.

This progressive reduction and convergence to similar estimates across measures (Table S8) capturing distinct social dimensions indicate that the apparent genetic signal in SFOH measures primarily reflects population stratification rather than trait-specific genetic architectures. If these measures captured genuine genetic effects on different social factors—neighborhood environment, loneliness, stress, discrimination—we would expect their heritability estimates to diverge after controlling for global ancestry structure. Instead, their convergence to a similar baseline suggests a single underlying confounder. The small residual heritability observed even with seven genetic PCs most likely represents incomplete control for fine-scale population structure, though genuine but minimal genetic effects on social factors or technical artifacts from survey data structure cannot be ruled out. This pattern of decrease and convergence was recapitulated when using PCA to summarize one-hot encoded SFOH data (Fig. S11), confirming that it is not an artifact of MCA but a robust property of the social data structure.

#### 2.4.2 The SSI demonstrates convergence without affecting trait heritability

The SSI demonstrates an important pattern. While it shows apparent heritability that, like the SFOH MC axes, decreases with genetic PC adjustment, it does not reduce trait heritability estimates when included as a covariate (Fig. S12). This distinction is informative. The SSI captures overall social similarity across all survey dimensions, and its correlation with genetic PCs is comparable to individual MC axes. Yet unlike specific MC axes, it does not affect trait heritability. This pattern suggests that correlation with population structure is necessary but not sufficient for SFOH measures to influence trait heritability estimates. Rather, measures must capture specific social dimensions that structure trait variation to act as effective covariates. The SSI result therefore supports the interpretation that reductions in trait heritability reflect reduction of genuine social confounding aligned with genetic structure rather than an artifact of including ancestry-correlated covariates.

### 2.5 SFOH measures and fundamental demographics strongly influence trait variance

We applied double generalized linear models to investigate how SFOH measures and demographics influence the dispersion of anthropometric and metabolic traits. All models included comprehensive adjustment for age, sex, race, and ethnicity while testing the two distinct SFOH measures: 20 SFOH MCs, and the SSI.

#### 2.5.1 Comprehensive dispersion effects across SFOH measures

All SFOH measures showed widespread significant effects on trait dispersion after multiple testing correction. Multiple MC axes showed widespread significant effects on trait dispersion (Fig. 5). In particular, SFOH MC axes demonstrate strong dispersion effects on anthropometric measures and cardiovascular traits. MCs 1 and 2 show broad significance, particularly on metabolic traits. Many MC axes reached nominal significance and a substantial amount survived stringent test correction.

**Fig. 5:**
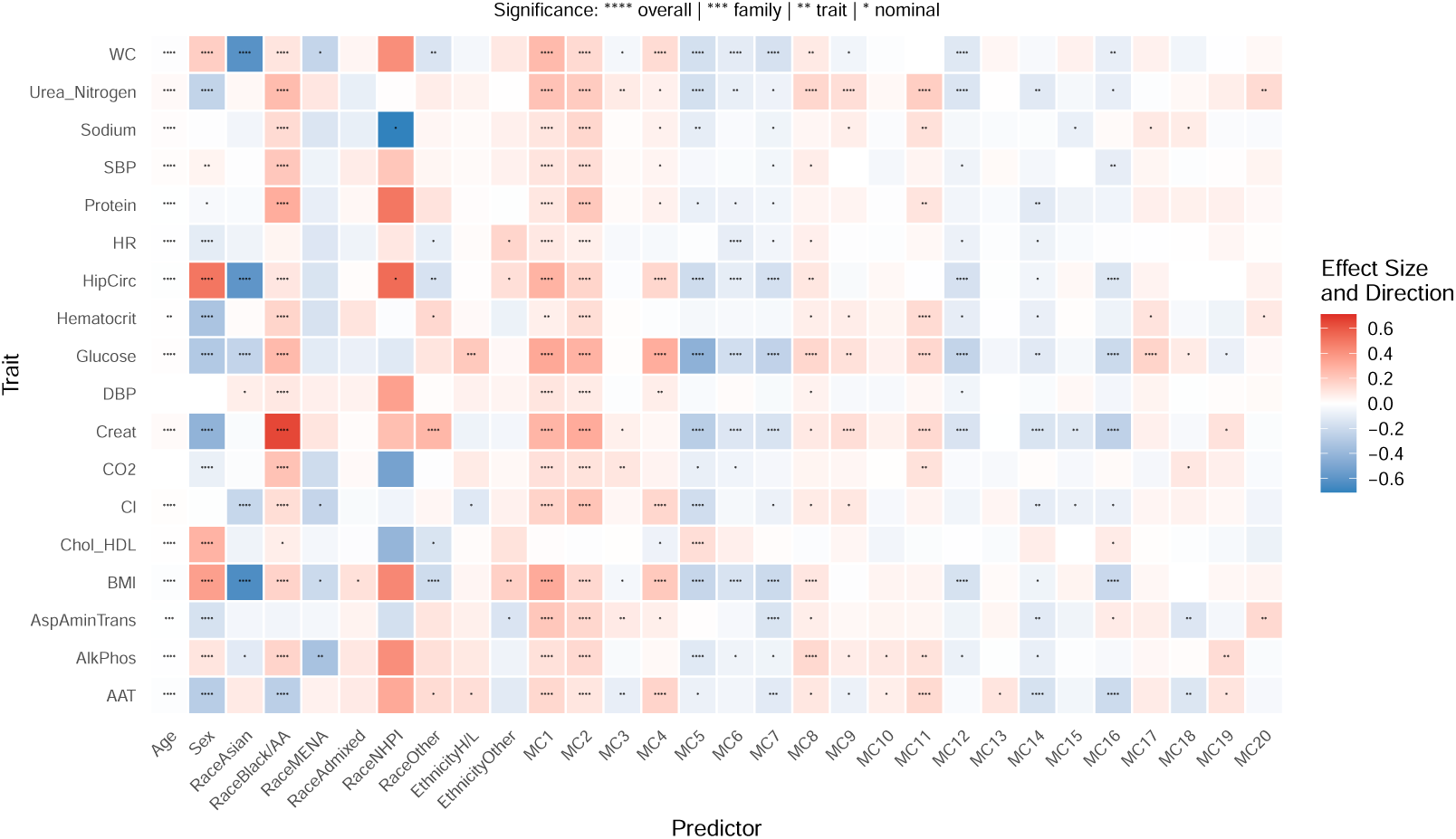
SFOH MC axes dispersion effects. Heatmap of significant predictors in trait (y-axis) dispersion models with demographics and SFOH MC axes (x-axis). The color gradient (blue to red) and intensity indicates the direction and magnitude of the effect size of the measure on the variance of each trait. Significance is based on a hierarchical Bonferroni correction with four tiers of stringency. Each test was assigned a significance level based on the most stringent threshold it passed and is indicated with “*”.

The SSI also demonstrated significant dispersion effects on traits (Fig. S13). Anthropometric measures, particularly hip and waist circumference show the strongest and most consistent association with the SSI. The SSI consistently reached overall significance after stringent test correction.

The two SFOH measurements: MC axes and SSI suggest that SFOH influence trait variance through both multi-dimensional approaches and a broader single summary. The convergence of both measures on anthropometric traits reinforces the robust connection between SFOH and body composition variance independent of demographic factors.

#### 2.5.2 Comprehensive dispersion effects of demographics

The demographic covariates we considered—age, sex, race, and ethnicity— contributed significantly to trait dispersion, independent of SFOH measures. Age demonstrated the most widespread influence across traits, suggesting that variance patterns systematically change with age. Sex, race, and ethnicity showed more targeted, but still significant, dispersion effect, indicating that demographic factors contribute to trait variance beyond their associations with SFOH measures. Further, demographic factors overall showed a greater magnitude of effect sizes, indicating that they are highly influential in trait variance provided they remain significant. The persistence of significant demographic factors after comprehensive SFOH adjustment underscores the importance of including these covariates to isolate true SFOH measure influences. Furthermore, the pattern of positive and negative effects across demographic covariates suggests that different groups may have distinct contributions to trait variance, highlighting the multidimensional nature of how demographics and SFOH shape trait variance.

### 2.6 Assessing the association between SFOH and genetic population structure

We assessed the extent to which SFOH and genetic population structure are associated by examining bidirectional relationships between SFOH MCs and genetic PCs in the combined analysis group.

#### 2.6.1 SFOH measures and genetic PCs show bidirectional but limited associations in the combined analysis group

First, to assess how social factors explain genetic structure, we regressed each of genetic PCs 1:10 on SFOH derived MC axes 1:40. A linear combination of MC axes explained non-trivial variation in genetic PC1 (*R*^2^ = 0.206), likely reflecting alignment between global ancestry and social groupings (Table S9). The variance explained by MC axes for genetic PCs 2:10 was considerably lower (max *R*^2^ for any genetic PC is 0.043).

Next, to assess how genetic structure explains social factor patterns, we regressed each of MC axes 1:20 on genetic PCs 1:10. Genetic PCs explained varying amounts of variation in MC structure. All twenty MCs showed statistically significant associations with genetic PCs, with *R*^2^ values ranging from 0.001 (MC12) to 0.076 (MC3) (Table S10). Several MCs demonstrated notable strong genetic associations, including the aforementioned MC3, MC4 (*R*^2^ = 0.047), MC1 (*R*^2^ = 0.044), and MC16 (*R*^2^ = 0.035). The consistent statistical significance across all components suggests broad genetic influence on the representation of SFOH measures. These axes likely capture SFOH topics such as feelings of healthcare and everyday discrimination, perceived stress, and neighborhood environment (Fig. 6).

**Fig. 6:**
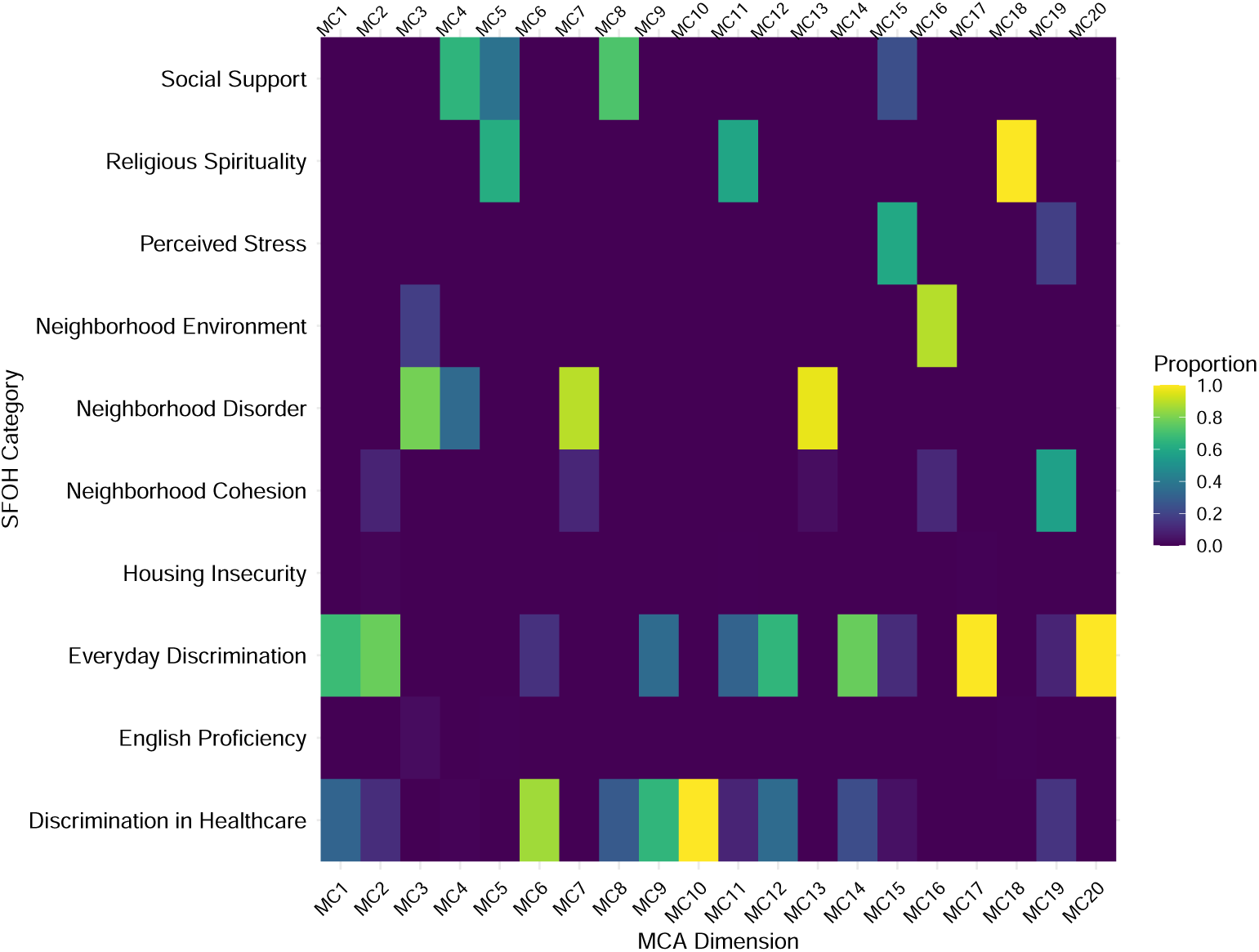
SFOH MCA dimension topic loadings heatmap. This heatmap depicts the contribution of each survey topic (y-axis) to SFOH MC axes (x-axis) derived from the All of Us Social Determinants of Health Survey. The color gradient (purple to yellow) represents each topic’s proportional weight, indicating its relative importance in defining each component. Each column sums to 1.

#### 2.6.2 Genetic PCs and SFOH measures constitute largely distinct sources of variation

These findings suggest that while SFOH MC axes and genetic PCs are statistically associated—particularly where social and ancestral structures align—the practical overlap is limited. Most of the variation in each domain does not covary with the other. This underscores the importance of treating social and genetic factors as distinct sources of variation in analyses of complex traits and reinforces the need to measure environmental exposures directly, rather than relying on race as a proxy. Further, this apparent distinction between domains supports the use of both SFOH MC axes and genetic PCs in downstream analyses.

## 3 Methods

### 3.1 Data source and Cohort Selection

We analyzed data from All of Us v7. To characterize the genetic ancestry structure of this diverse cohort, we performed PCA and present the population centroids for PC1 and PC2 (Fig. S14). For a detailed visualization of the genetic PC structure of the All of Us dataset, we refer readers to Figures 1 and 2 in [28]. The dataset includes over one billion genetic variants and has facilitated the identification of genetic associations with various diseases.

We included individuals who had genetic data available, completed the Social Determinants of Health survey, self-identified male or female sex at birth, and who had reported a self-identified racial identity from The Basics survey. To maximize sample size, we included individuals identifying as “Hispanic or Latino”. The final combined analysis group comprised 85,963 individuals.

As a sensitivity analysis, we repeated key analyses in a subset of 48,146 individuals of European ancestry who additionally identified as “White” and “Not Hispanic or Latino” which we refer to as the “European-ancestry subgroup”. We determined genetic ancestry using the All of Us genetic similarity categories from the auxiliary file ancestry preds.tsv, which group individuals based on their genetic similarity to reference populations from the Genome Aggregation Database (gno-mAD), Human Genome Diversity Project (HGDP), and 1000 Genomes. Specifically, we used the “1KGP-HGDP-EUR-like (EUR or European) category to define the European-ancestry subgroup.

### 3.2 Quality Control

Genotyping was performed using a custom Illumina array built on the design frame-work of the Global Screening Array. The genome build used is hg38/GRCh38. Please refer to the All of Us Research Program technical documentation and data release notes for further details. We conducted variant filtering using PLINK 2.0 [29] on the All of Us version 7 controlled-tier array data, restricted to autosomal SNPs. This reassigned variant identifiers and removed all duplicate variants. To mitigate the effects of linkage disequilibrium, we pruned variants using a sliding window approach (200 variants per window, with a step size of 50 variants) and an *r*^2^ threshold of 0.2. We excluded variants with a minor allele frequency below 0.01, a genotype missingness rate exceeding 1%, deviating from Hardy-Weinberg equilibrium (*p <* 1*x*10^−6^), and individuals with more than 2% missing genotype calls. We assessed relatedness up to the third degree using KING [30] (kinship coefficient ≥ 0.0625), and removed one member from each related pair. After quality control, we retained 285,123 unrelated individuals and 56,961 high-quality common variants for genetic PC analysis (Fig. S2).

### 3.3 Trait Processing

We selected 18 clinically relevant anthropometric and metabolic traits known to demonstrate variation across self-identified racial groups, enabling us to investigate how additional social factors might contribute to these patterns. These traits include: alanine aminotransferase (AAT), alkaline phosphatase (AlkPhos), aspartate aminotransferase (AspAminTrans), body mass index (BMI), carbon dioxide (CO2), chloride (Cl), cholesterol in HDL (CholHDL), creatinine (Creat), diastolic blood pressure (DBP), glucose, heart rate (HR), hematocrit, hip circumference (Hip-Circ), protein, sodium, systolic blood pressure (SBP), urea nitrogen (UreaNitro), and waist circumference (WC).

Following trait selection, we performed data cleaning and deduplication procedures prior to downstream analyses. For individuals with multiple measurements, we retained the most recent value. To mitigate the influence of extreme values, we calculated the IQR for each biomarker independently, and observations falling outside these bounds were truncated to the corresponding threshold values. Systolic blood pressure was adjusted for antihypertensive medication using a binary indicator variable. Sample sizes varied by trait due to differential measurement availability across participants.

### 3.4 Estimating Variance Contribution by Genome-wide Complex Trait Analysis (GCTA)

We estimated *h*^2^ using HE regression as implemented in the Genome-wide Complex Trait Analysis software (GCTA). For our primary inference, we report HE-CP point estimates from HE regression and standard errors (SE) calculated by jackknifing over individuals. 95% confidence intervals (CIs) were constructed as *HE* − *CP* ± 1.96(*SE*). To evaluate whether SFOH measures alter heritability estimates, and whether SFOH measures themselves may be heritable, we compared the 95% CIs between pairs of nested models. Non-overlapping CIs were interpreted as evidence of a significant difference in heritability estimates between models, which provides a conservative test, as this criterion is more stringent than a direct significance test at *α*=0.05.

Additionally, we computed z-scores to quantify the magnitude of change in heritability estimates between models, calculated as:

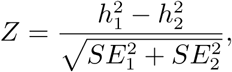

where 1 and 2 represent heritability estimates and respective SEs from the two models being contrasted.

### 3.5 Estimating Dispersion Components with Double Generalized Models

We employed double generalized linear models (dglms) to investigate the effect of SFOH measures on both the mean and the variance of traits. Dglms extend traditional generalized linear models by simultaneously modeling both the mean and the dispersion components of the response variable, allowing us to test whether the variance of the trait differs across levels of predictor variables. We implemented the dglm package in R with a Gaussian distribution for the mean model and the default log link for the variance model. We fit the following mean and variance models for 18 traits:

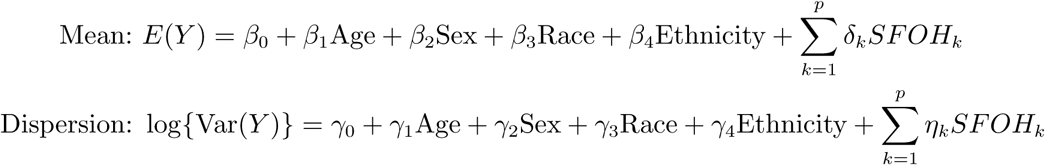

where *SFOH_k_* represents the *k*th component of one of two SFOH measures analyzed in separate models: the first 20 MC axes (*p* = 20) or the SSI as a single measure (*p* = 1). To account for multiple comparisons, we applied a hierarchical Bonferroni correction with four tiers of stringency. Each test was assigned a significance level based on the most stringent threshold it passed:

1. **Overall**: p *<* 0.05 / (total number of tests across all traits and SFOH measures)
2. **Family-wise**: p *<* 0.05 / (number of tests within the SFOH measure)
3. **Trait-wise**: p *<* 0.05 / (number of predictors per trait)
4. **Nominal**: p *<* 0.05 (uncorrected)

We then labeled predictors according to the highest significance level achieved (overall, family-wise, trait-wise, or nominal).

### 3.6 Social Determinants of Health Survey and SFOH Derivation

We derived Social factors of health (SFOH) measures from the All of Us Social Determinants of Health (SDOH) survey (v7), which captures data on factors such as income, education, and housing stability. Our analysis used the survey data as it was available in the v7 dataset. Out of the 397,732 participants eligible for the survey by June 30, 2022, 117,783 (29.6%) had completed it. From these respondents, 94,030 had genotyping array data available. We one-hot encoded the raw survey responses to create a structured dataset for downstream analysis. From this encoded data, we generated two distinct SFOH measures: multiple correspondence analysis axes (MCA), and a Social Similarity Index (SSI), as detailed below.

#### 3.6.1 Survey Preparation

Survey items were organized into themes (i.e. conceptual domains grouping related questions, such as “perceived stress,” “social support,” “loneliness,” etc.) based on the SFOH Task Force’s consensus-based conceptual frameworks. All available survey completers (n=117,783) were used to generate the SFOH measures to ensure stable estimates.

#### 3.6.2 MCA

To capture the multidimensional nature of SFOH, we applied dimensionality reduction to one-hot encoded survey data. Our primary analysis used MCA, implemented in Python using the prince package, as it is the methodologically appropriate technique for categorical data. We merged the resulting component scores into a single analysis dataset using unique subject identifiers. We include up to the first 40 SFOH MC axes, which capture the major axes of social-factor heterogeneity in our cohort, as linear covariates in downstream analyses.

As a sensitivity analysis to assess the robustness of our conclusions to the choice of dimension reduction technique, we also performed PCA on the same one-hot encoded survey data using prcomp in R. Results from PCA-based analyses were qualitatively consistent with the primary MCA findings throughout. Key PCA analyses are presented in the Supplement, and referenced in the main text where relevant.

#### 3.6.3 Environmental Relatedness Matrix (ERM)

We constructed a environmental relatedness matrix (ERM) from the same one-hot encoded SFOH survey data. The ERM is conceptually analogous to a GRM, here we derive it entirely from SFOH survey data. In this approach, every individual’s binary response vector forms a row of the data matrix

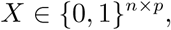

where *n* is the number of individuals and *p* is the total number of response-level columns. To ensure that each question contributes equally regardless of how many response levels it has, we form a diagonal weight matrix

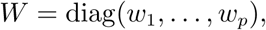

with

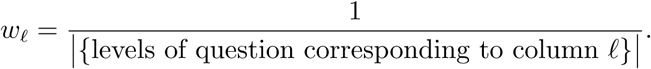

We define the weighted design matrix

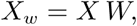

so that

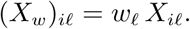

To account for missing responses, we define a binary mask matrix

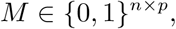

by setting

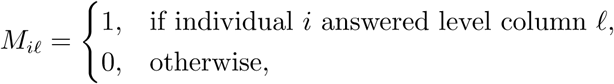

and its weighted version

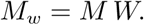

All matrix operations are carried out on the sparse weighted matrices *X_w_* and *M_w_*. Although ERM*_ii_* = 1 holds by definition, we never explicitly compute or store diagonal entries.

#### 3.6.4 Social Similarity Index (SSI)

The SSI provides an individual-level summary of how similar each person’s survey responses are to all other participants in the cohort, accounting for the structure of the survey instrument. Conceptually, individuals with more similar social circumstances (as captured by their survey responses) will have higher SSI values, reflecting shared social environments. The SSI is computed by comparing each person’s response pattern to every other person’s responses across all SFOH survey questions, with appropriate weighting to ensure that questions with more response options do not disproportionately influence the similarity calculation. Mathematically, this is accomplished through the ERM. Rather than computing or storing the full *n* × *n* ERM, we summarize each individual’s social similarity to the cohort using the ERM row sums. We define each row sum as

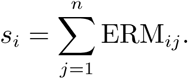

We compute the row sum in *O*(*n p*) time by first defining the two vectors

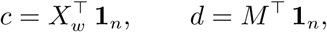

Here, *d* gives the unweighted sum of answered items per column. We then compute the individual-level weighted sums

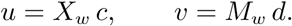

Each individual’s SSI is then

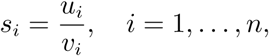

and we set any indeterminate 0*/*0 to zero (so that individuals with no overlapping responses are treated as unrelated). We standardize the SSI to have mean zero and unit variance. The standardized SSI captures each individual’s overall social similarity to the cohort which we include in downstream regression models to account for shared social context.

## 4 Discussion

Our results demonstrate that SFOH measures reduce HE regression heritability estimates for a subset of anthropometric and metabolic traits—particularly BMI, hip and waist circumference, and HDL cholesterol—in this multi-ancestry analysis group.

HE regression’s sensitivity to SFOH adjustment demonstrates how social factors may proxy genetic structure, leading to inflated estimates when unmodeled. This makes HE regression a valuable tool for detecting potential confounding. SFOH-associated variance detected by HE regression may represent confounding or indirect genetic effects rather than direct additive genetic variance.

The heritability of SFOH measures themselves provides direct evidence that they capture population structure. Several SFOH measures appeared moderately heritable when adjusting for basic demographics or a limited number of genetic PCs (Fig. S7 and Fig. S8). However, with comprehensive adjustment using seven genetic PCs, these heritability estimates decreased dramatically (Table S8). Measures that remained significantly different than zero converged to a common low baseline (Table S7). This pattern was recapitulated in the European-ancestry subgroup with the recalculated SFOH MCs (Fig. S9), indicating that SFOH measures capture stratification that persists within more ancestrally homogeneous populations. If different SFOH measures reflected distinct genetic influences on different social factors, we would expect divergent estimates after PC adjustment; instead, their convergence indicates they all capture the same underlying confounder. This finding explains why including SFOH measures as covariates reduces trait heritability estimates—they remove stratified variance that would otherwise be misattributed to genetics by HE regression.

This finding has important implications for genetic studies. First, it demonstrates that inadequate control for population structure can create spurious evidence of genetic effects on environmental and social factors. Second, it explains why including SFOH measures as covariates reduces trait heritability estimates—they capture confounding social stratification that would otherwise be misattributed to genetics. Third, it suggests that the small residual heritability (≈ 1 − 2%) observed even with seven PCs likely represents fine-scale population structure rather than meaningful genetic effects on social environments.

Furthermore, several SFOH measures that showed this pattern exhibit widespread, though small, effects on trait dispersion even after adjusting for age, sex, race, and ethnicity (Fig. 5), indicating they influence both trait means and variances. Notably, while the SSI showed a similar initial heritability pattern, it does not reduce trait heritability estimates, suggesting that reductions are driven by specific social dimensions rather than global social similarity.

The interpretation of the initial apparent heritability of SFOH measures is complex, as it may reflect genuine genetic influence on social factors through gene-environment correlation [31], residual population stratification not fully captured by the first three genetic PCs, technical artifacts from survey data correlation structure, social stratification [32], or environmental confounding [33]—processes that can induce genetic associations with social factors even without direct biological pathways. Our finding that these heritability estimates decrease with comprehensive genetic PC adjustment strongly suggests that, in our multi-ancestry cohort, residual population stratification is the primary driver. Regardless of mechanism, these SFOH measures introduce structured variance that can inflate trait heritability estimates in HE regression when unmodeled. While adjusting for heritable covariates risks removing true genetic signal [34], our results indicate that in this context, SFOH adjustment primarily removes stratification that would otherwise be misattributed to genetics.

We acknowledge several methodological limitations. First, the comprehensive dispersion models include up to 33 predictors and may capture sample-specific patterns that do not generalize; these results should be viewed as hypothesis-generating rather than definitive. Second, while our multi-ancestry design limits direct generalization to any single population group, it provides advantages for detecting population stratification. The pronounced genetic structure in multi-ancestry cohorts makes SFOH-ancestry correlations more detectable, suggesting our findings may represent a conservative bound on confounding that could be subtler but still present in more homogeneous populations. Third, while our use of seven genetic PCs was sufficient to demonstrate that SFOH heritability estimates decrease, even this number may not capture all fine-scale population structure. Residual stratification could therefore still be partially captured by SFOH measures or influence trait heritability estimates. Fourth, our use of self-identified race—a social construct that imperfectly captures both genetic ancestry and social experiences—risks conflating genetic background with socially mediated SFOH exposures; between group comparisons are not interpretable as true population differences and are indistinguishable from statistical artifacts in this study.

These results highlight that SFOH structure can meaningfully alter SNP-based heritability estimates and trait variance patterns. A small number of social dimensions—social support, neighborhood environment, feelings of discrimination, stress, and spirituality—appear especially important in shaping the variance of complex traits. The convergence of SFOH heritability estimates across diverse social dimensions provides evidence that these measures primarily capture population stratification rather than independent genetic effects on social environments. This work underscores that for traits with strong social patterning, particularly anthropometric measures, measuring and adjusting for specific social environmental factors may be necessary to avoid misattributing social influences to genetic effects. Future work should focus on methods to disentangle the roles of genetic structure, social environments, and their interactions in shaping trait variance.

## 5 Data availability

Individual-level genotype and survey data used in this study are available to registered researchers through the All of Us Researcher Workbench (https://workbench. researchallofus.org/) following Data and Research Center (DRC) access procedures. Genomic data (v7 array release) and survey responses, including the Social Determinants of Health survey, are part of the All of Us [35] controlled tier dataset. Other information is available upon request.

## Acknowledgments

We thank the All of Us Research Program and its participants. We thank Lily Hamilton for thoughtful feedback that led to the improvement of this manuscript. This research was supported by NHGRI CEGS RM1: RM1HG010461 (NSA) and NIH NCI U54CA267738 (SAM), The Cornell University Graduate School Deans Excellence Fellowship (OSRA), and The Cornell University Graduate School Walter Schonlenk and Leister Fellowships (JLL).

## Declarations

The authors declare no competing interests.

## Supplemental Material

**Fig. S1:**
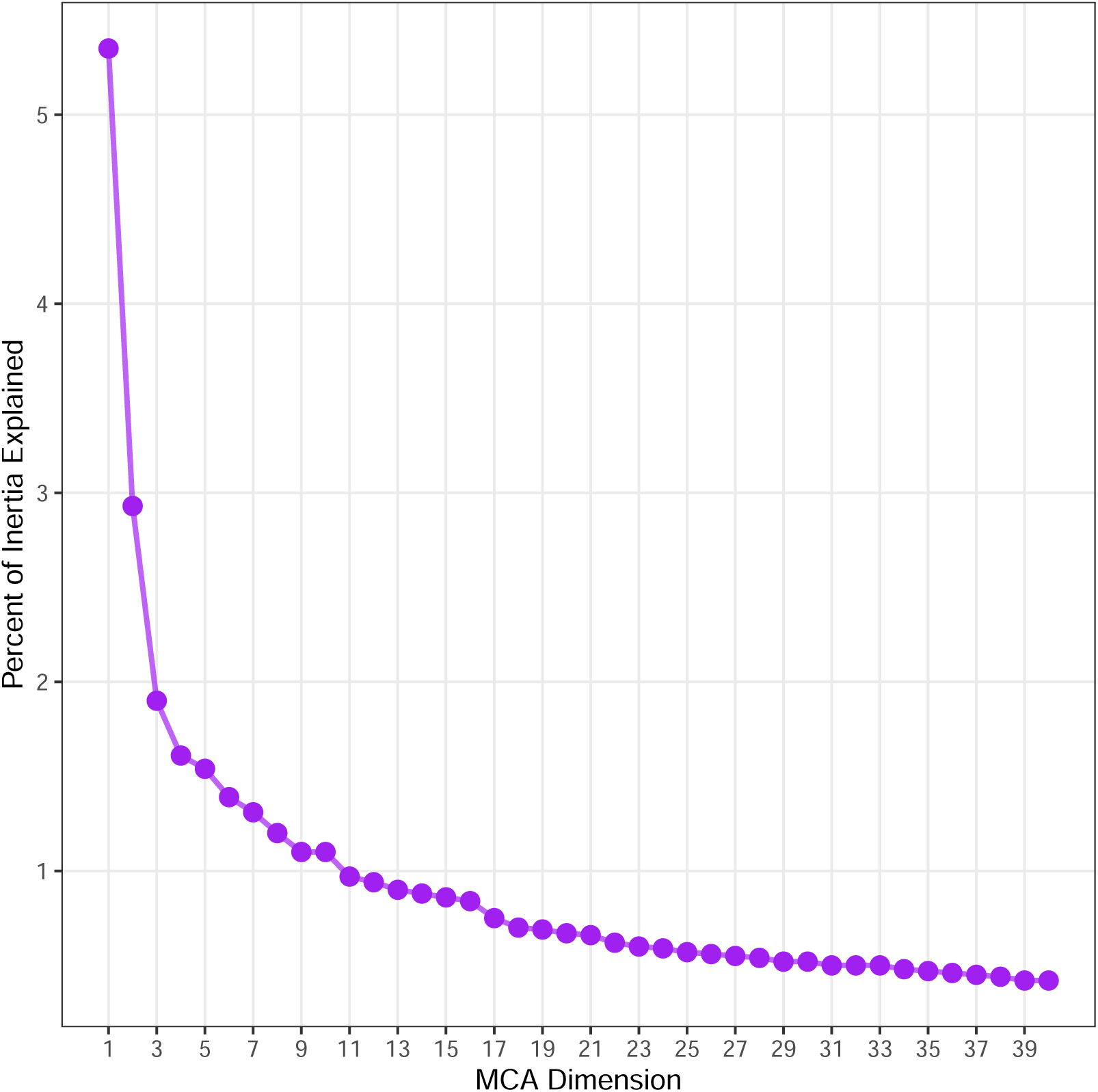
SFOH MCA scree plot. The lead measures, SFOH MC axes 1:6, account for 14.73% of variance. Comprehensive measures, SFOH MC axes 1:40, account for 38.02% of variance.

**Fig. S2:**
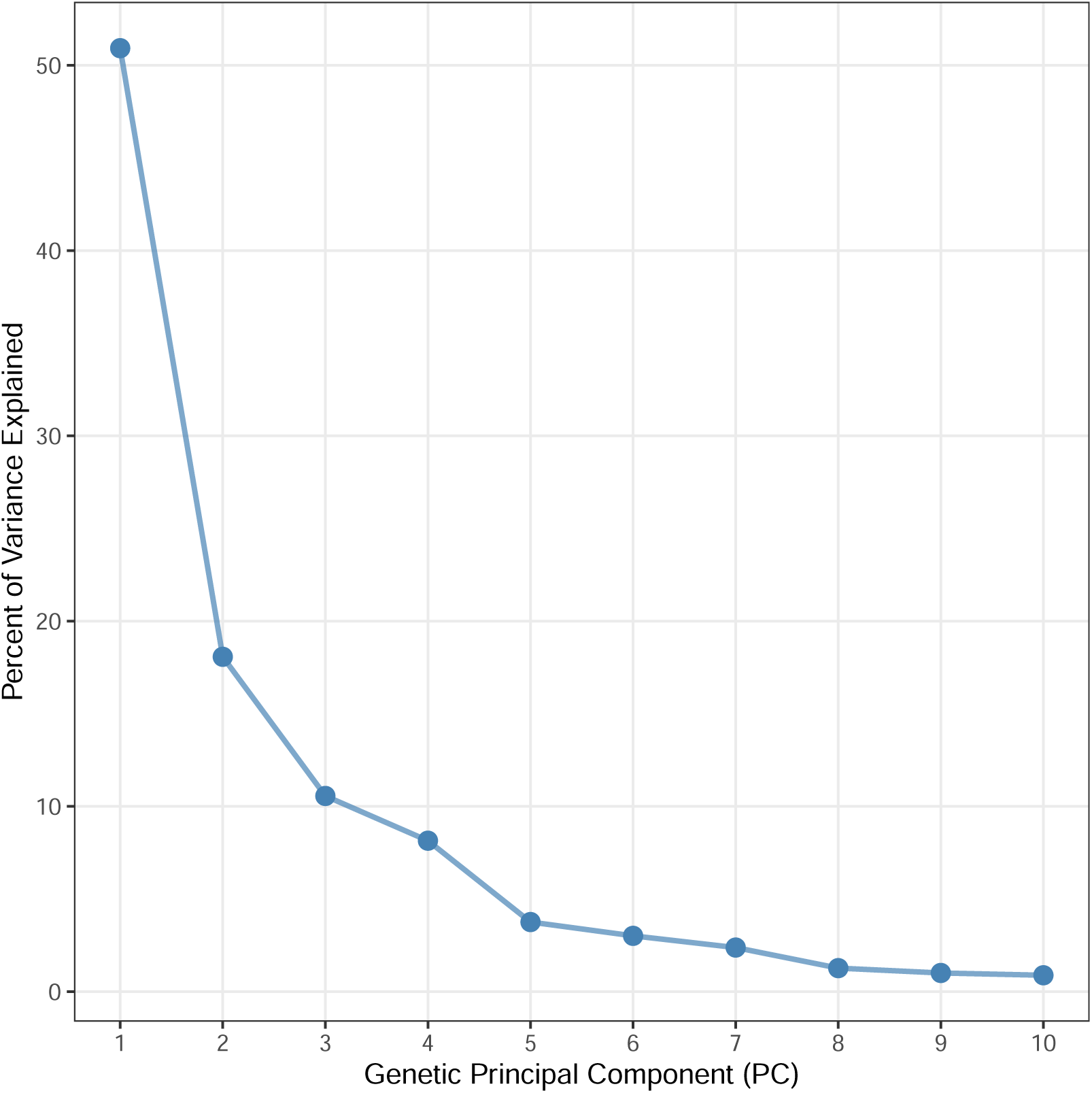
Genetic PCA scree plot. Genetic PCs 1:3 account for 79.56% of variance, while genetic PCs 1:7 account for 96.84% of variance.

**Fig. S3:**
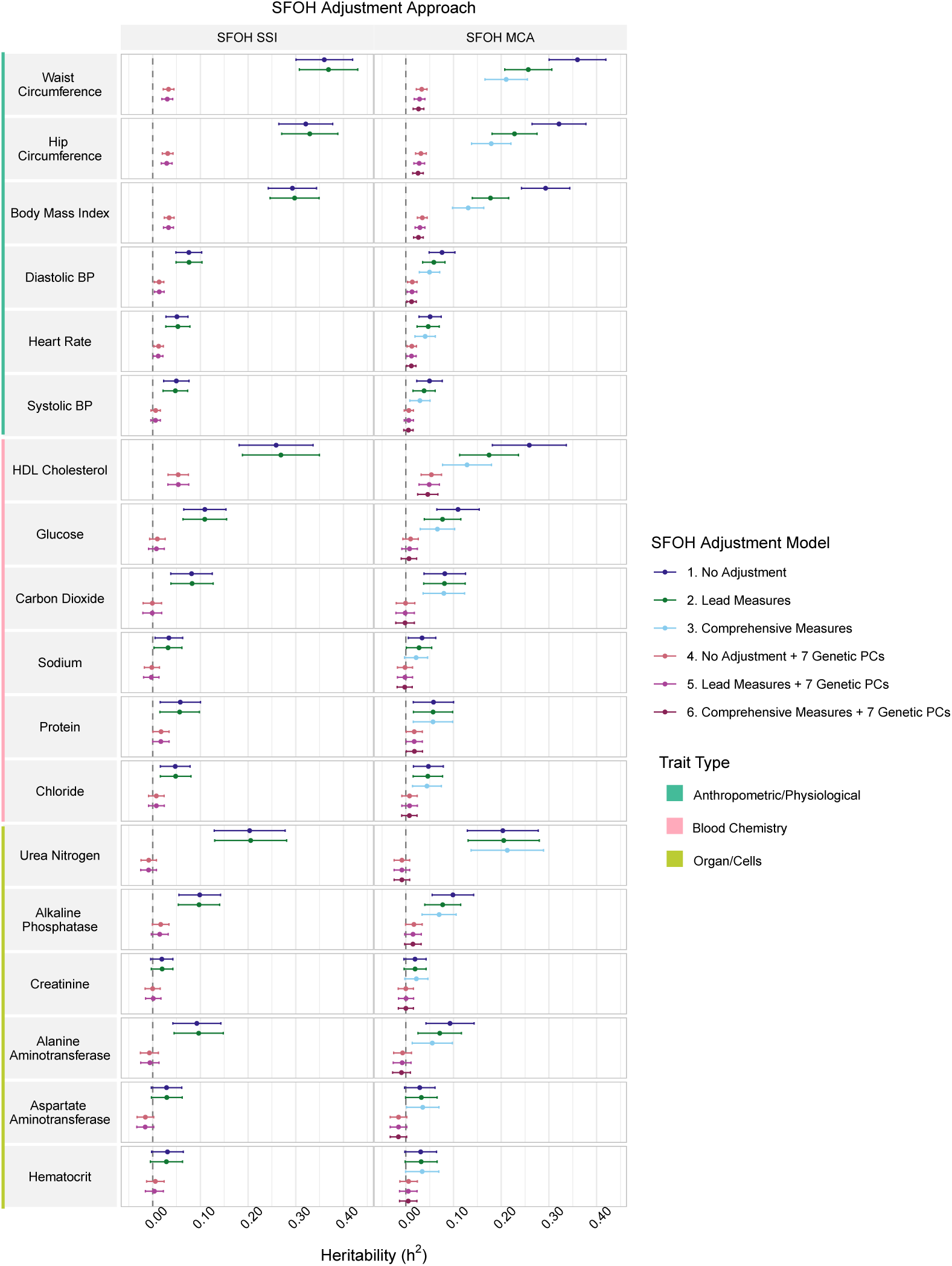
Female Haseman-Elston (HE) regression across SFOH-covariate models. HE regression heritability estimates of female participants with 95% confidence intervals for 18 traits (rows), calculated using two Social Factors of Health (SFOH) summarization strategies (columns). All models are adjusted for age, race, and ethnicity. Models with SFOH adjustment include lead (MCs 1:6 or SSI) or comprehensive (MCs 1:40) adjustment. Patterns are consistent with the combined analysis group (Fig. 2). Points represent HE-CP estimates; intervals are 95% CIs from jackknife standard errors. The dashed line indicates a heritability of zero.

**Fig. S4:**
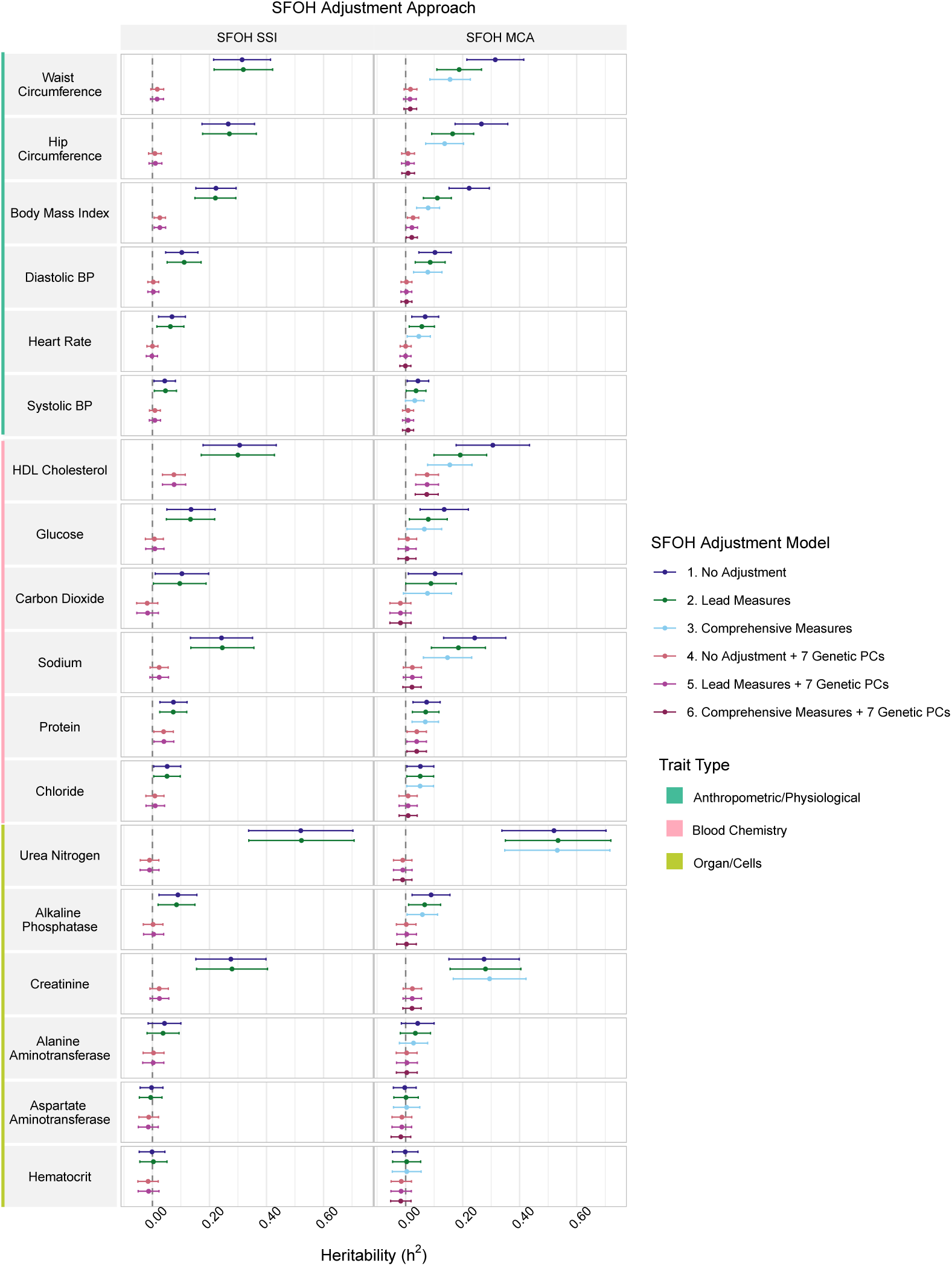
Male Haseman-Elston (HE) regression across SFOH-covariate models. HE regression heritability estimates of male participants with 95% confidence intervals for 18 traits (rows), calculated using two Social Factors of Health (SFOH) summarization strategies (columns). All models are adjusted for age, race, and ethnicity. Models with SFOH adjustment include lead (MCs 1:6 or SSI) or comprehensive (MCs 1:40) adjustment. Patterns are consistent with the combined analysis group (Fig. 2). Points represent HE-CP estimates; intervals are 95% CIs from jackknife standard errors. The dashed line indicates a heritability of zero.

**Fig. S5:**
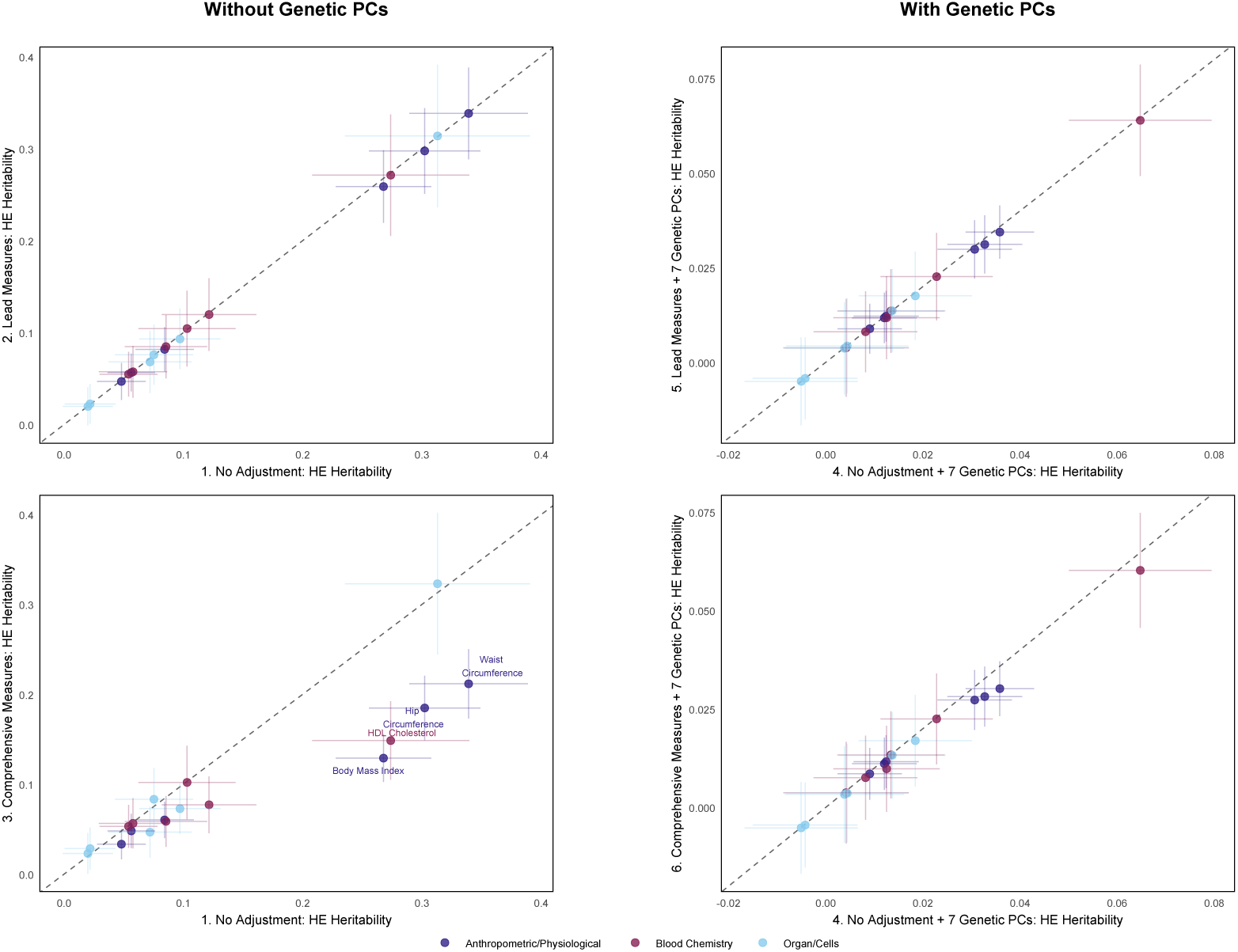
Combined analysis group Haseman-Elston (HE) regression across SFOH PC adjustment models. HE regression heritability estimates for all relevant participants with 95% confidence intervals for 18 traits, calculated using SFOH PCs. Panels are arranged in a 2×2 grid: models without genetic PCs (left) and with genetic PCs (right). Each panel shows a pairwise model comparison with model numbers noted at the beginning of the and y-axis labels. All models are adjusted for age, sex, race, and ethnicity. Models with SFOH adjustment include lead (PCs 1:3) or comprehensive (PCs 1:20) adjustment. Adding SFOH PCs (models 2,3) reduces heritability estimates for BMI, HDL cholesterol, hip circumference, and waist circumference relative to the base model (model 1). Points represent HE-CP estimates; intervals are 95% CIs from jackknife standard errors. The dashed line indicates the y = x line. Results are broadly consistent with MCA findings, though MCA is the statistically appropriate method for categorical data.

**Fig. S6:**
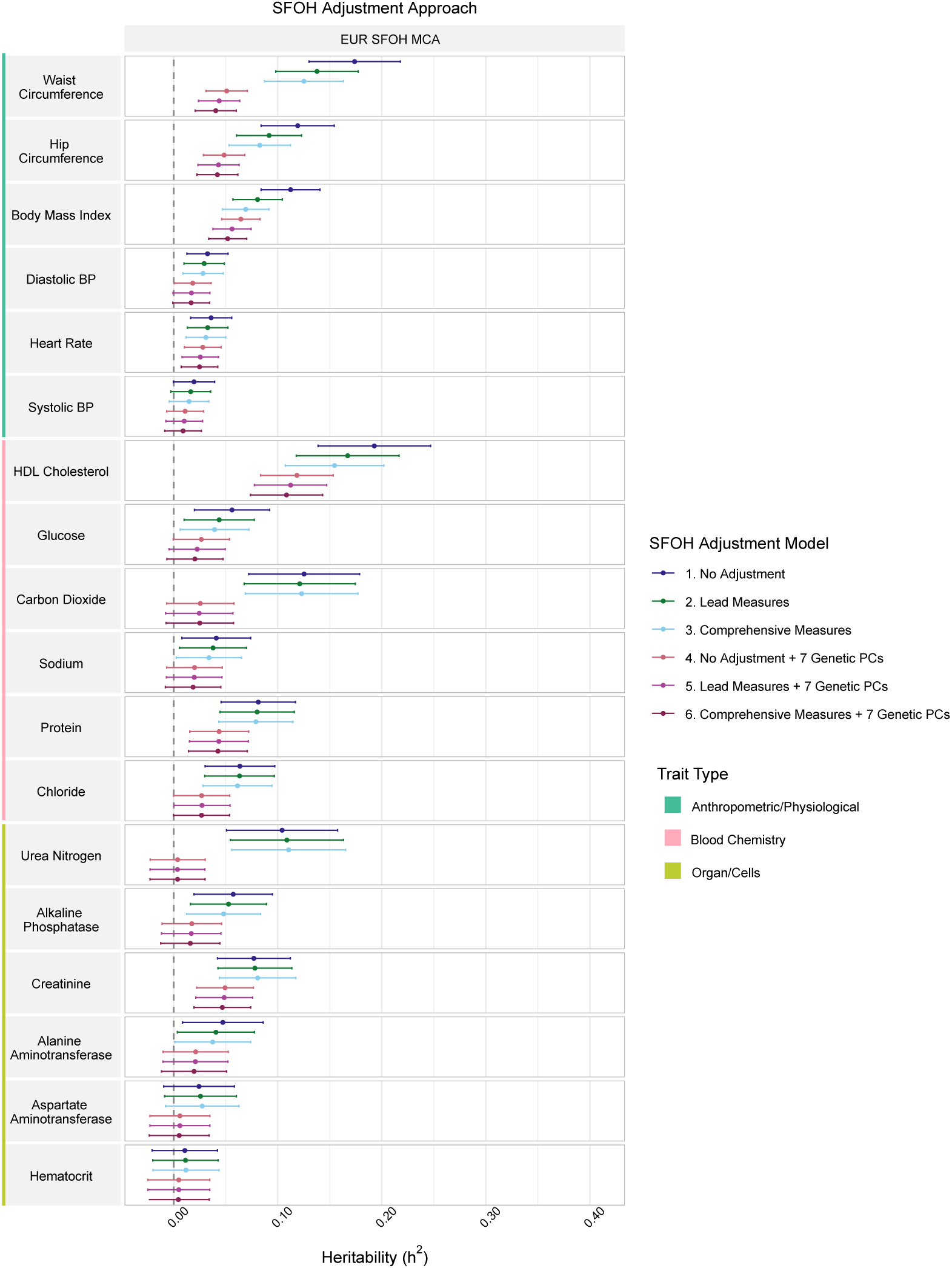
European-ancestry subgroup Haseman-Elston (HE) regression across SFOH MC adjustment models. HE regression heritability estimates for the European-ancestry subgroup of individuals with 95% confidence intervals for 18 traits (rows), calculated using SFOH MCs caluclated from this strict subset of individuals. All models are adjusted for age and sex. Models with SFOH adjustment include lead (strict MCs 1:6) or comprehensive (strict MCs 1:40) adjustment. Patterns are consistent with the combined analysis group (Fig. 2). Points represent HE-CP estimates; intervals are 95% CIs from jack-knife standard errors. The dashed line indicates a heritability of zero.

**Fig. S7:**
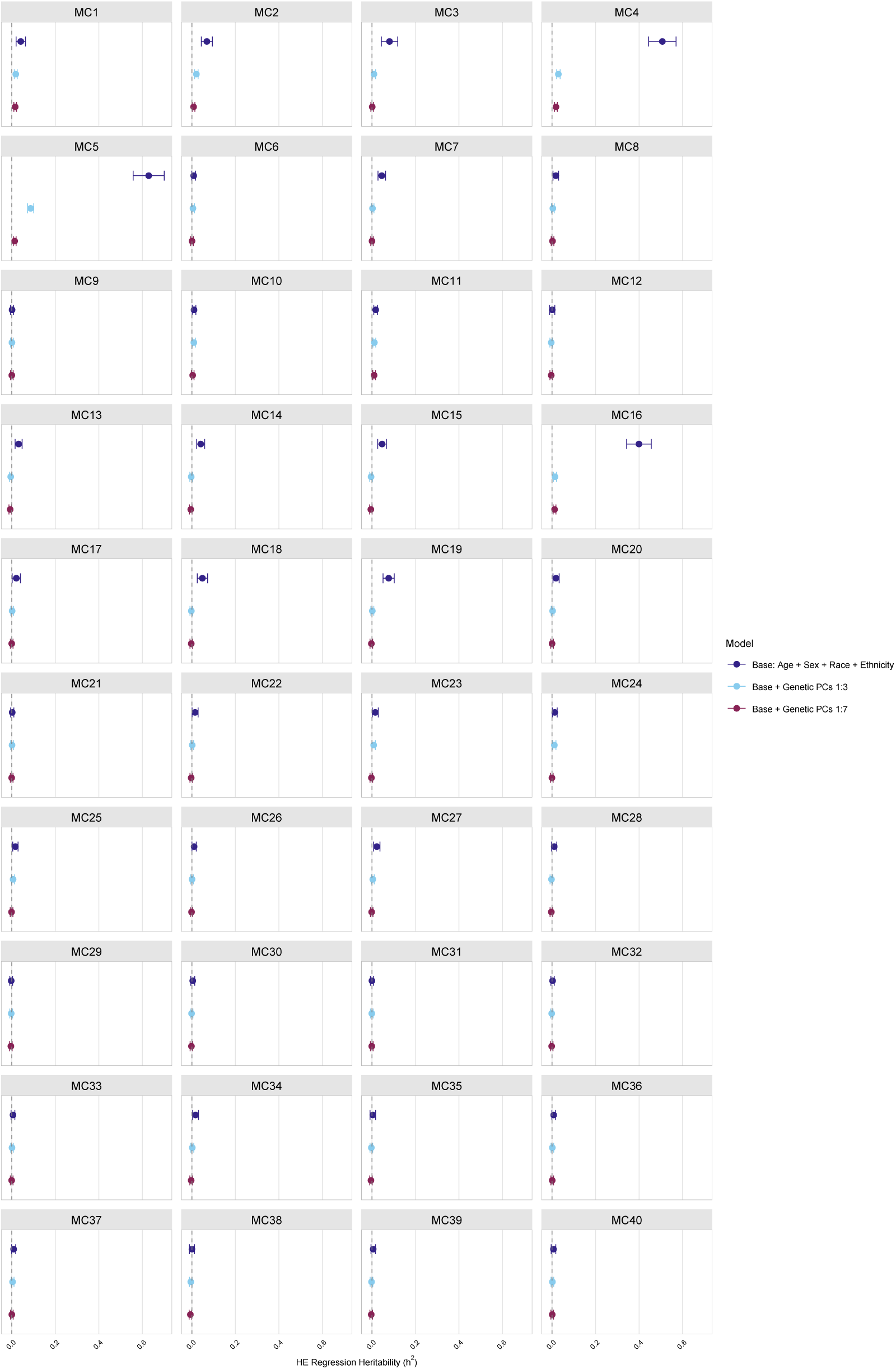
Combined analysis group Haseman-Elston (HE) regression heritability of SFOH MC axes. HE regression heritability estimates for all relevant participants with 95% confidence intervals for 40 SFOH MC Axes calculated using models with progressive genetic PC adjustment. Points represent HE-CP estimates; intervals are 95% CIs from jackknife standard errors. The dashed line indicates a heritability of zero. HE regression heritability estimates demonstrate dramatic reductions and convergence with genetic PC adjustment indicating these measures capture population stratification.

**Fig. S8:**
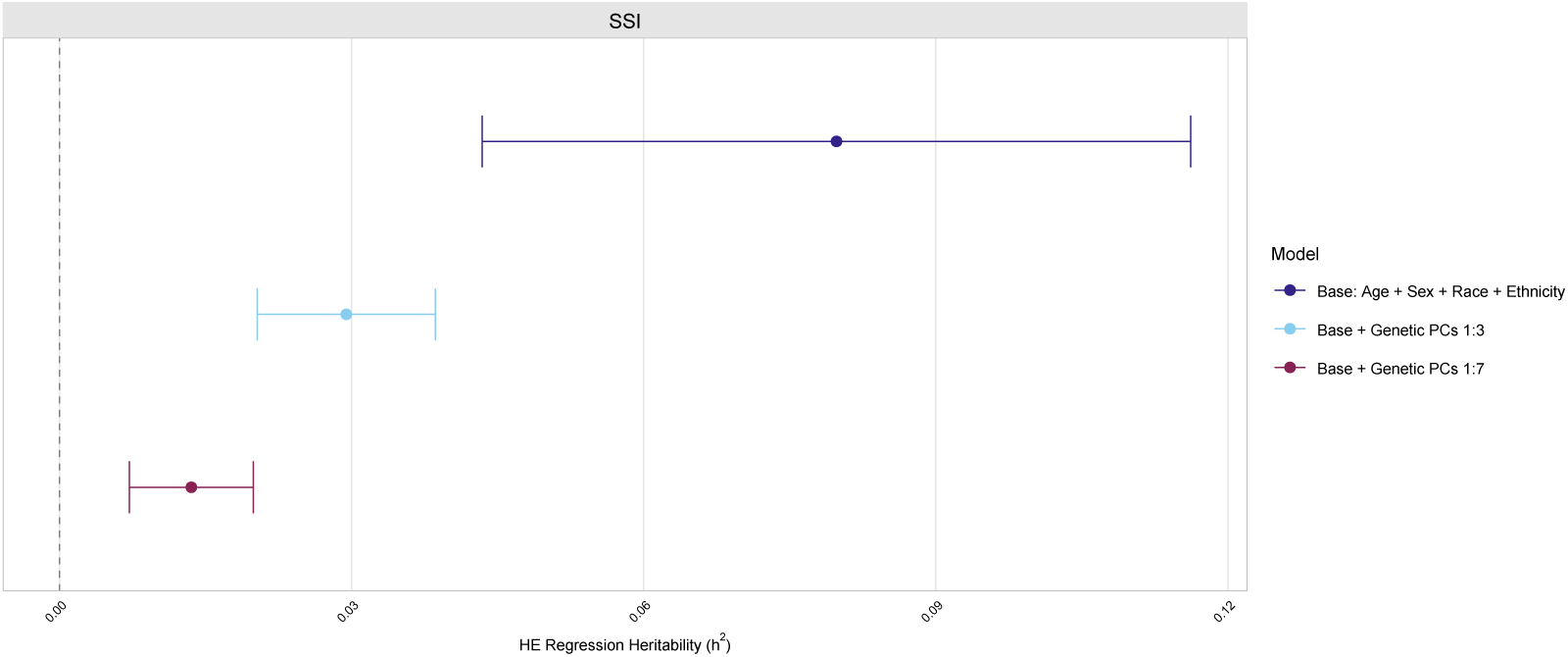
Combined analysis group Haseman-Elston (HE) regression heritability of the SSI. HE regression heritability estimates for all relevant participants with 95% confidence intervals for the SSI calculated using three nested models: demographic adjustment only, demographics and the top three genetic PCs, and demographics and the top seven genetic PCs. Points represent HE-CP estimates; intervals are 95% CIs from jackknife standard errors. The dashed line indicates a heritability of zero. HE regression heritability estimates show progressive reduction with genetic PC adjustment. While significant at all adjustment levels, the SSI heritability decreases from *h*^2^ = 0.080 (base) to 0.029 (3 PCs) to 0.014 (7 PCs), demonstrating that apparent heritability primarily reflects population stratification.

**Fig. S9:**
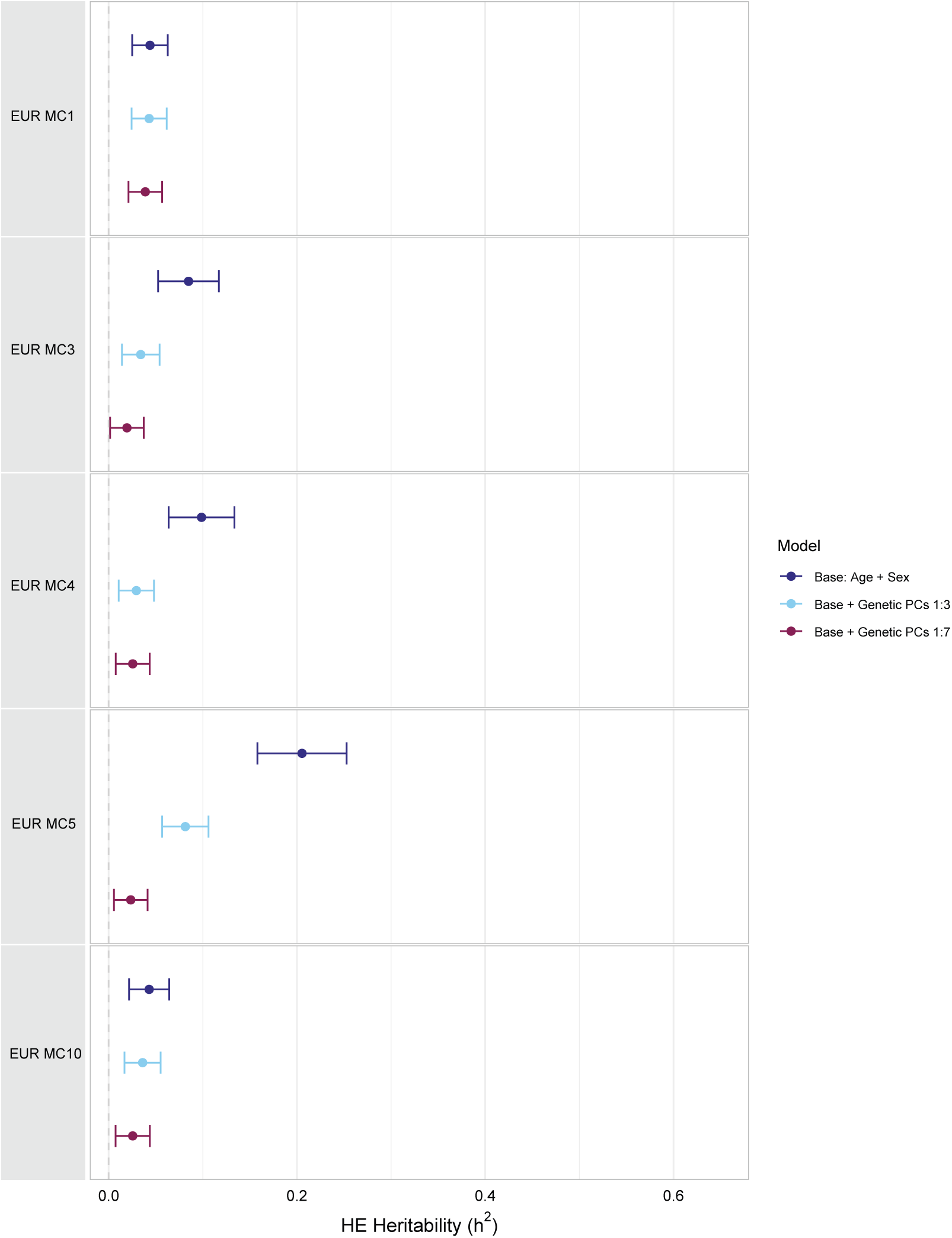
European-ancestry subgroup Haseman-Elston (HE) regression heritability of SFOH MCs measures calculated in this strict subset that remain heritable after including seven genetic PCs. HE regression heritability estimates for the European-ancestry subgroup of individuals with 95% confidence intervals for SFOH measures that appear heritable after comprehensive genetic PC adjustment. Points represent HE-CP estimates; intervals are 95% CIs from jackknife standard errors. The dashed line indicates a heritability of zero. HE regression heritability estimates show progressive reduction with genetic PC adjustment, demonstrating that apparent heritability primarily reflects population stratification.

**Fig. S10:**
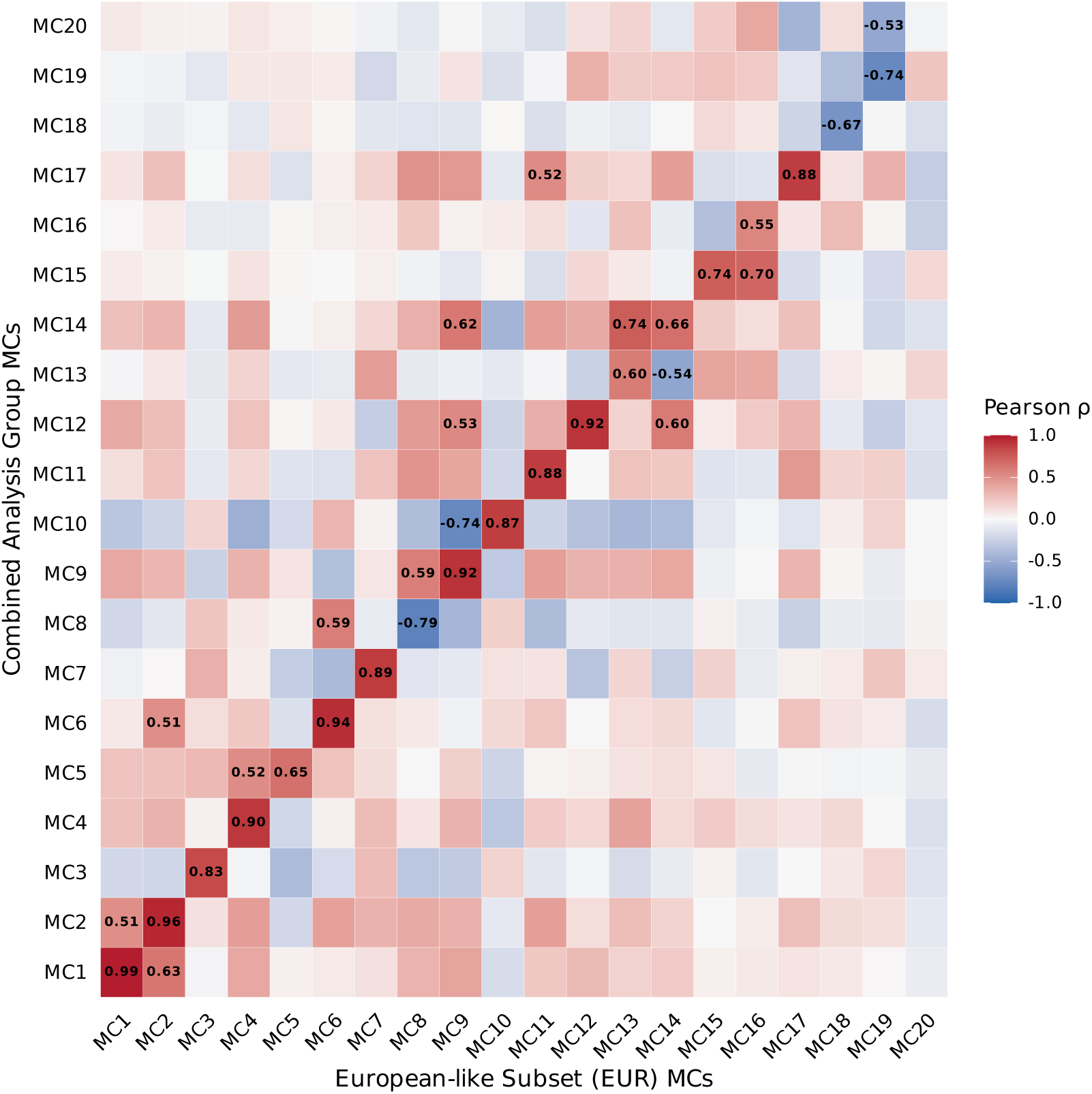
Heatmap of Pearson correlations between SFOH MCA dimensions from two population subsets. Columns represent the first 20 SFOH MCs derived from the combined analysis group; rows represent corresponding dimensions from the European-ancestry subgroup. The strong correlations on the diagonal indicate that primary latent structures are conserved. Deviations from the diagonal suggest some population-specific variation in some dimensions.

**Fig. S11:**
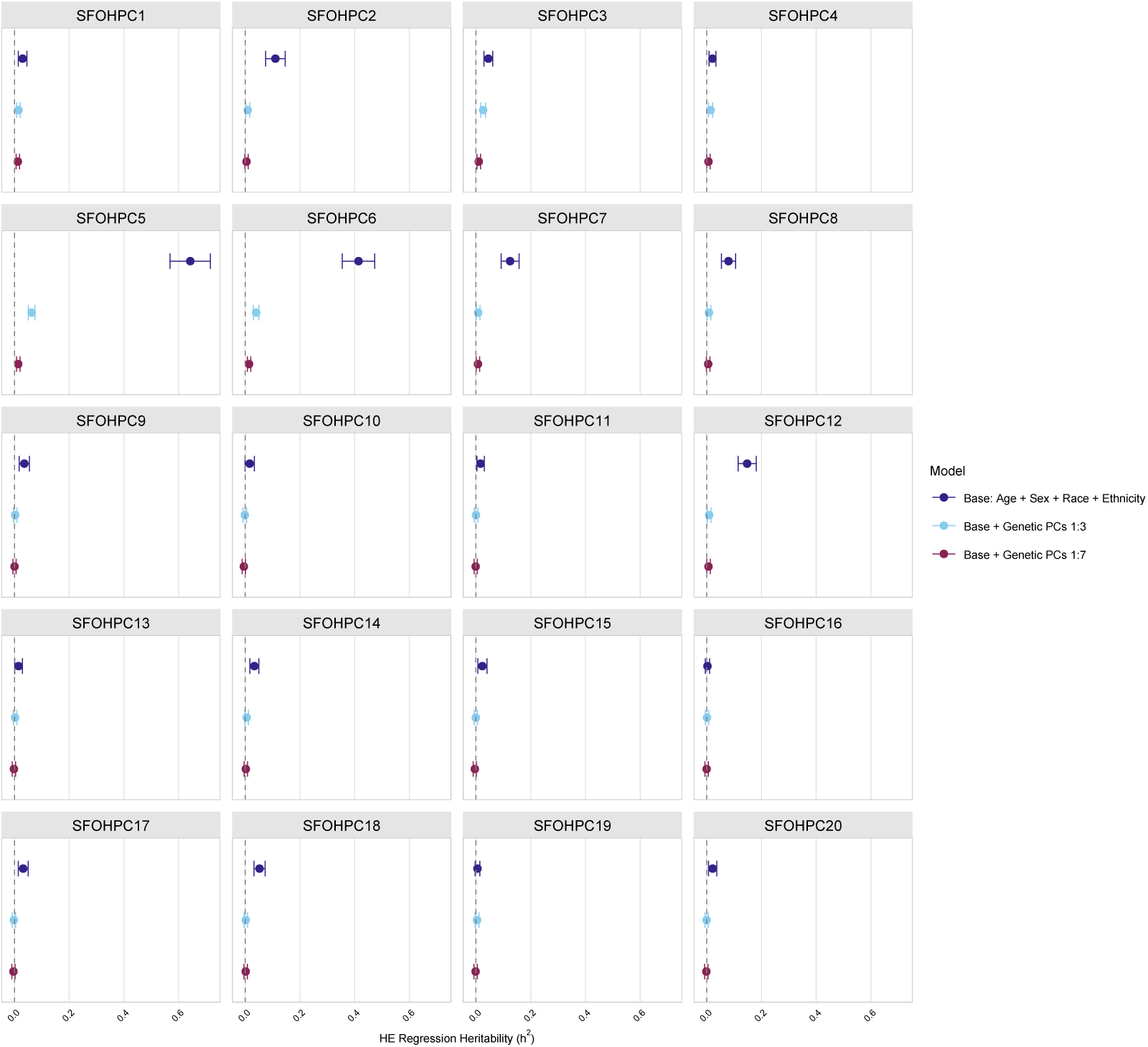
Combined analysis group Haseman-Elston (HE) regression heritability of SFOH PCs. HE regression heritability estimates for all relevant participants with 95% confidence intervals for 20 SFOH PCs calculated using models with progressive genetic PC adjustment. Points represent HE-CP estimates; intervals are 95% CIs from jackknife standard errors. The dashed line indicates a heritability of zero. HE regression heritability estimates demonstrate dramatic reductions and convergence with genetic PC adjustment indicating these measures capture population stratification.

**Fig. S12:**
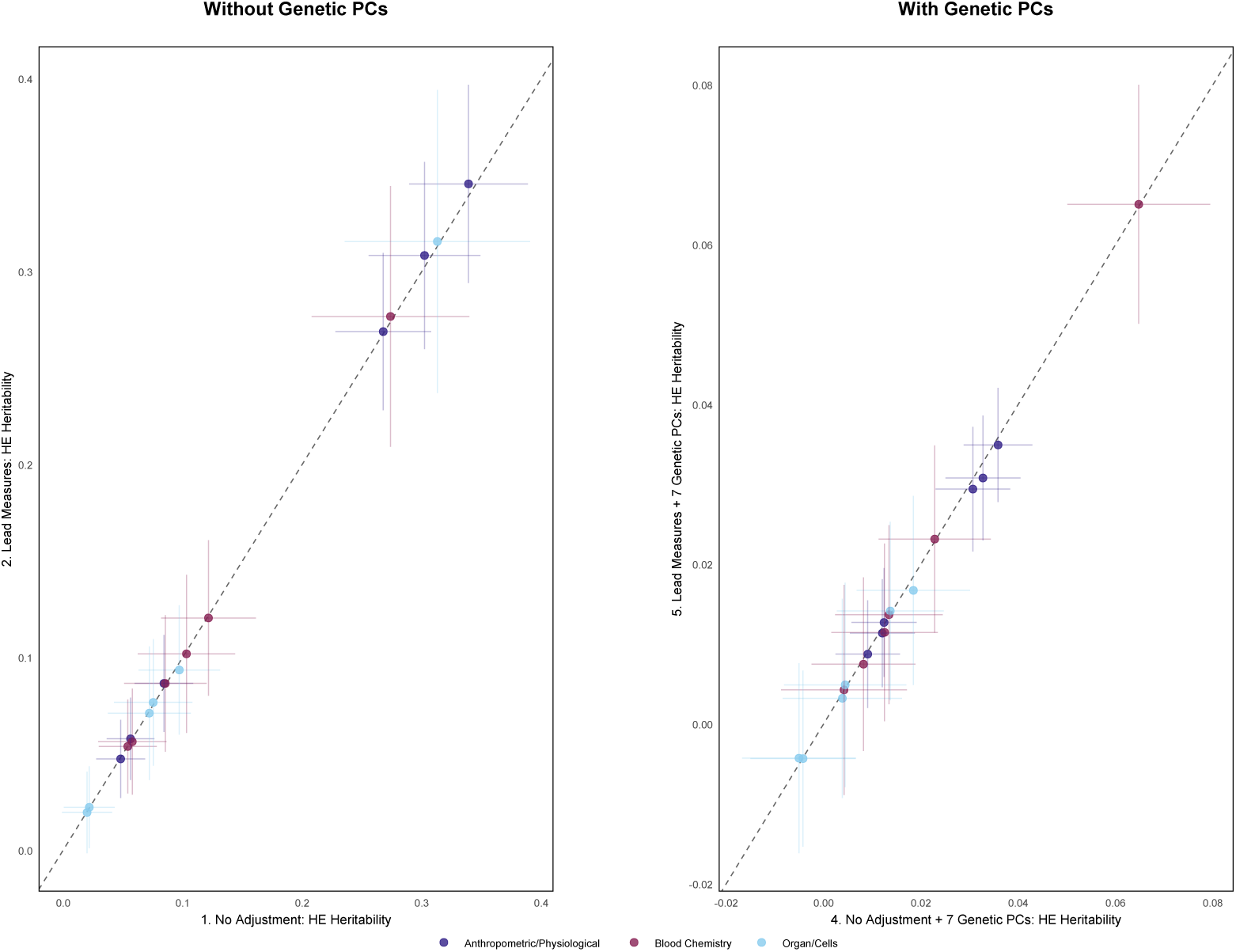
Combined analysis group Haseman-Elston (HE) regression across SFOH SSI adjustment models. HE regression heritability estimates for all relevant participants with 95% confidence intervals for 18 traits, calculated using SFOH SSI. Models without genetic PCs (left) and with genetic PCs (right). Each panel shows a pairwise model comparison with model numbers noted at the beginning of the x- and y-axis labels. All models are adjusted for age, sex, race, and ethnicity. Adding SFOH SSI (models 2,5) does not reduce heritability estimates relative to the base model (model 1). Points represent HE-CP estimates; intervals are 95% CIs from jackknife standard errors. The dashed line indicates the y = x line.

**Fig. S13:**
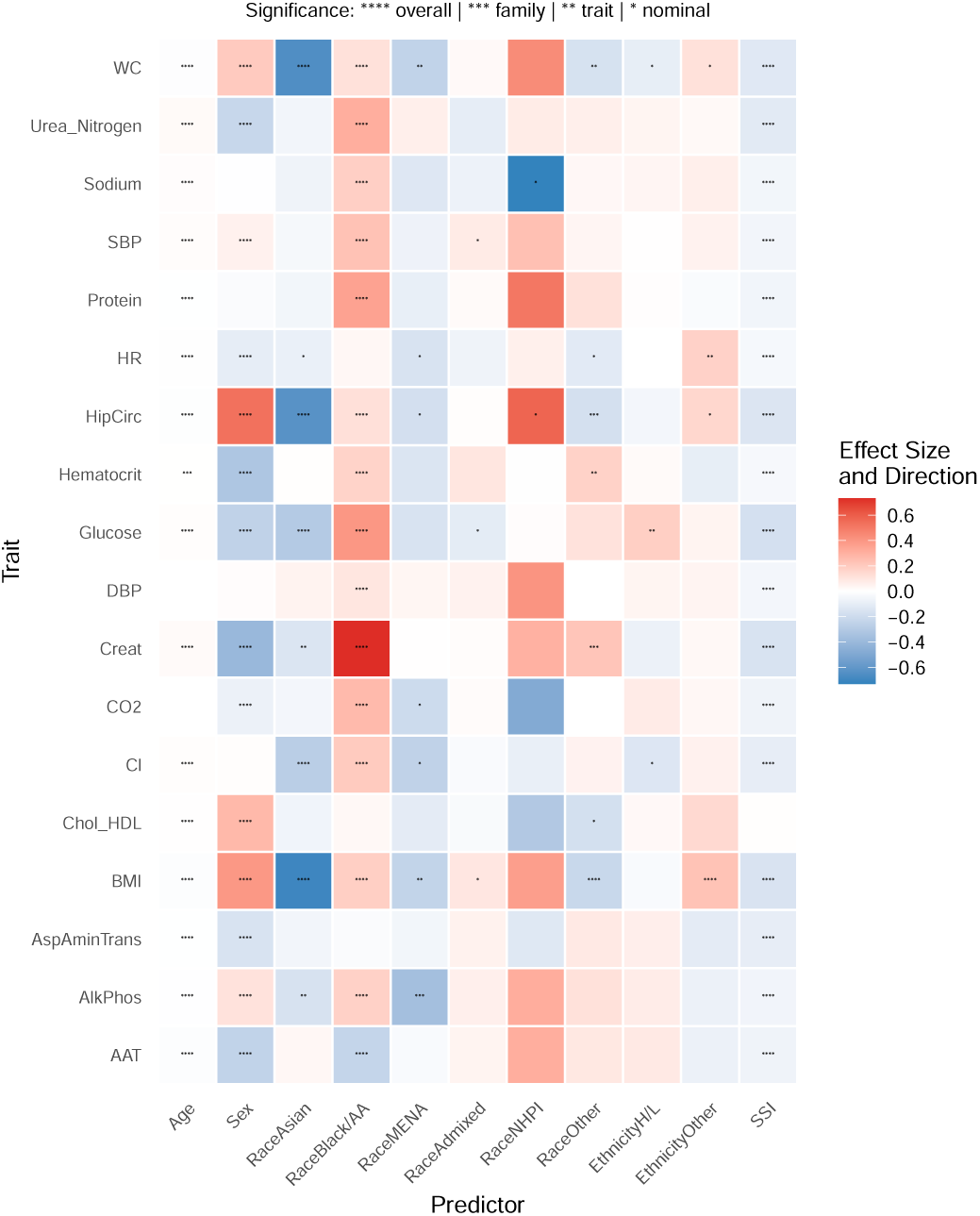
SFOH SSI dispersion effects. Heatmap of significant predictors in trait (y-axis) dispersion models with demographics and SSI (x-axis). The color gradient (blue to red) and intensity indicates the direction and magnitude of the effect size of the measure on the variance of each trait. Significance is based on a hierarchical Bonferroni correction with four tiers of stringency. Each test was assigned a significance level based on the most stringent threshold it passed and is indicated with “*”.

**Fig. S14:**
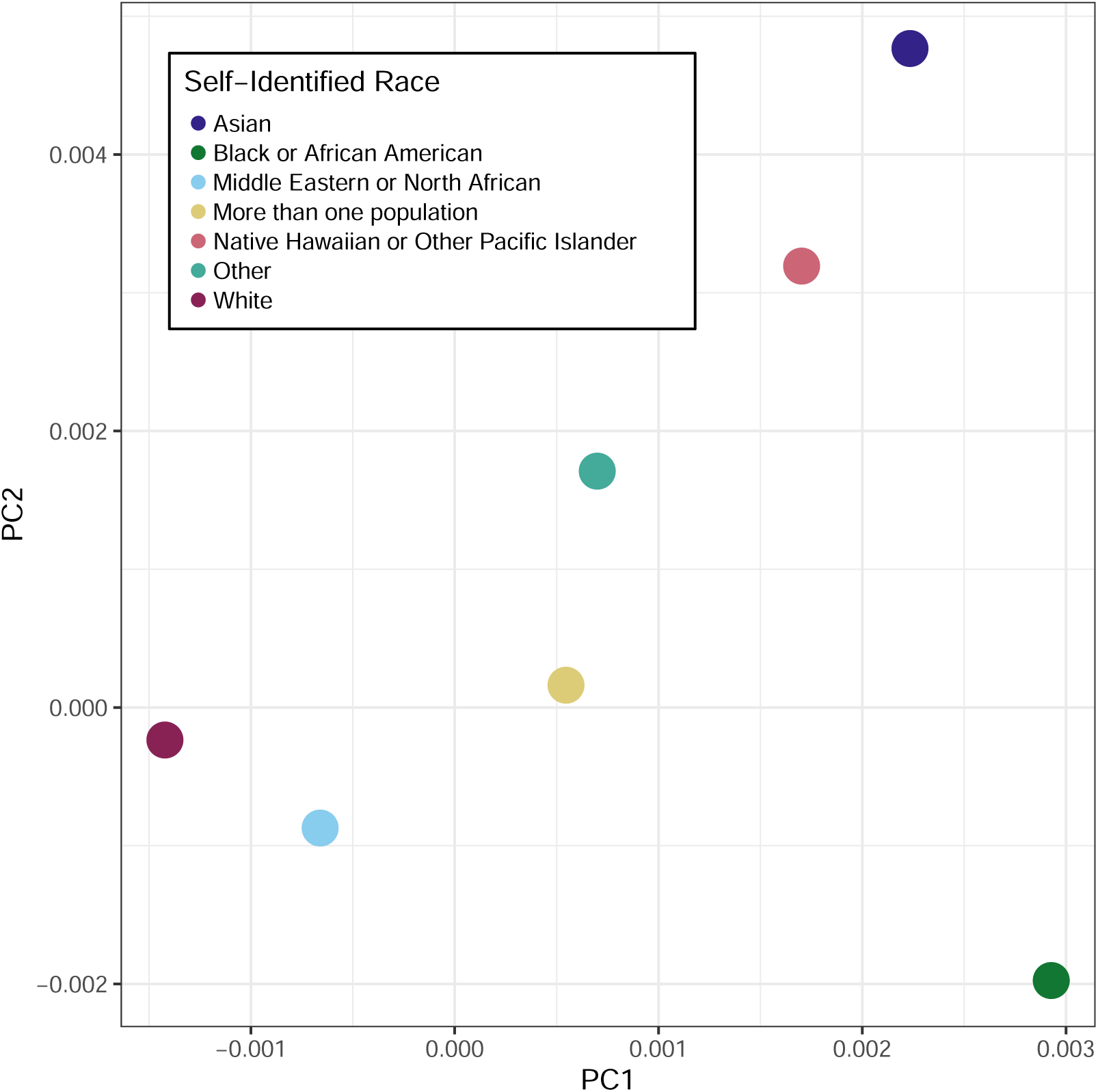
Population centroids from PCA of the All of Us genetic data. Points represent mean PC1 (x-axis) and PC2 (y-axis) values for each self-identified racial group, summarizing the overall genetic ancestry structure.

**Table S1:** Sample size of females and males that have genetic data and responded to the SFOH survey.

|  | **Sex** | **Count** |
| --- | --- | --- |
| 1 | Female | 54967 |
| 2 | Male | 30996 |

**Table S2:** Sample size of each self-identified race group that has genetic data and responded to the SFOH survey.

| **Self-Identified Race** | **Count** |
| --- | --- |
| Black or African American | 6513 |
| Asian | 2318 |
| Middle Eastern or North African | 380 |
| More than one population | 1511 |
| Native Hawaiian or Other Pacific Islander | 42 |
| White | 67164 |
| Other | 8035 |

**Table S3:** Overall Missingness by Theme.

| **Theme** | **Total Responses** | **Missing** | **% Missing** |
| --- | --- | --- | --- |
| Religious Attendance | 88016 | 10716 | 12.2 |
| Neighborhood Environment | 704128 | 51475 | 7.3 |
| Housing Instability | 88016 | 4594 | 5.2 |
| Perceived Stress | 880160 | 37119 | 4.2 |
| Neighborhood Disorder | 1144208 | 34943 | 3.1 |
| Housing Insecurity | 88016 | 2641 | 3.0 |
| Discrimination in Healthcare Settings | 616112 | 17083 | 2.8 |
| Everyday Discrimination | 792144 | 20631 | 2.6 |
| Social Support | 704128 | 15945 | 2.3 |
| Neighborhood Cohesion | 352064 | 7227 | 2.1 |
| Loneliness | 704128 | 14207 | 2.0 |
| Religious Spirituality | 528096 | 8518 | 1.6 |
| Food Insecurity | 176032 | 1622 | 0.9 |

**Table S4:** Percentage of question level non-response by “Theme” and Self-Identified Race. Rows containing less than 20 individuals removed as per All of Us data usage agreement.

| Theme | Race | Total Responses | Missing (%) |
| --- | --- | --- | --- |
| Discrimination in Healthcare | Black or African American | 46081 | 2079 (4.5%) |
| Discrimination in Healthcare | Other | 67312 | 2909 (4.3%) |
| Discrimination in Healthcare | Middle Eastern or North African | 2681 | 88 (3.3%) |
| Discrimination in Healthcare | White | 472815 | 11513 (2.4%) |
| Discrimination in Healthcare | Asian | 16261 | 299 (1.8%) |
| Discrimination in Healthcare | More than one population | 10654 | 187 (1.8%) |
| Everyday Discrimination | Black or African American | 59247 | 2621 (4.4%) |
| Everyday Discrimination | Other | 86544 | 3538 (4.1%) |
| Everyday Discrimination | White | 607905 | 13792 (2.3%) |
| Everyday Discrimination | Asian | 20907 | 383 (1.8%) |
| Everyday Discrimination | More than one population | 13698 | 223 (1.6%) |
| Everyday Discrimination | Middle Eastern or North African | 3447 | 55 (1.6%) |
| Food Insecurity | Black or African American | 13166 | 274 (2.1%) |
| Food Insecurity | Other | 19232 | 339 (1.8%) |
| Food Insecurity | Asian | 4646 | 52 (1.1%) |
| Food Insecurity | White | 135090 | 927 (0.7%) |
| Housing Insecurity | Black or African American | 6583 | 387 (5.9%) |
| Housing Insecurity | Other | 9616 | 518 (5.4%) |
| Housing Insecurity | Asian | 2323 | 60 (2.6%) |
| Housing Insecurity | White | 67545 | 1625 (2.4%) |
| Housing Insecurity | More than one population | 1522 | 29 (1.9%) |
| Housing Instability | White | 67545 | 3668 (5.4%) |
| Housing Instability | Other | 9616 | 506 (5.3%) |
| Housing Instability | Black or African American | 6583 | 259 (3.9%) |
| Housing Instability | Asian | 2323 | 90 (3.9%) |
| Housing Instability | More than one population | 1522 | 48 (3.2%) |
| Loneliness | Black or African American | 52664 | 1865 (3.5%) |
| Loneliness | Other | 76928 | 2698 (3.5%) |
| Loneliness | Middle Eastern or North African | 3064 | 53 (1.7%) |
| Loneliness | White | 540360 | 9128 (1.7%) |
| Loneliness | Asian | 18584 | 292 (1.6%) |
| Loneliness | More than one population | 12176 | 162 (1.3%) |
| Neighborhood Cohesion | Black or African American | 26332 | 833 (3.2%) |
| Neighborhood Cohesion | Other | 38464 | 1155 (3.0%) |
| Neighborhood Cohesion | White | 270180 | 5063 (1.9%) |
| Neighborhood Cohesion | Middle Eastern or North African | 1532 | 22 (1.4%) |
| Neighborhood Cohesion | More than one population | 6088 | 65 (1.1%) |
| Neighborhood Cohesion | Asian | 9292 | 81 (0.9%) |
| Neighborhood Disorder | Other | 125008 | 5532 (4.4%) |
| Neighborhood Disorder | Black or African American | 85579 | 3776 (4.4%) |
| Neighborhood Disorder | White | 878085 | 24622 (2.8%) |
| Neighborhood Disorder | Middle Eastern or North African | 4979 | 131 (2.6%) |
| Neighborhood Disorder | Asian | 30199 | 542 (1.8%) |
| Neighborhood Disorder | More than one population | 19786 | 319 (1.6%) |
| Neighborhood Environment | Black or African American | 52664 | 4794 (9.1%) |
| Neighborhood Environment | Other | 76928 | 6401 (8.3%) |
| Neighborhood Environment | Middle Eastern or North African | 3064 | 220 (7.2%) |
| Neighborhood Environment | White | 540360 | 38320 (7.1%) |
| Neighborhood Environment | More than one population | 12176 | 726 (6.0%) |
| Neighborhood Environment | Asian | 18584 | 982 (5.3%) |

Table S4 – continued from previous page
| Theme | Race | Total Responses | Missing (%) |
| --- | --- | --- | --- |
| Perceived Stress | Black or African American | 65830 | 3945 (6.0%) |
| Perceived Stress | Other | 96160 | 5316 (5.5%) |
| Perceived Stress | Middle Eastern or North African | 3830 | 173 (4.5%) |
| Perceived Stress | White | 675450 | 26568 (3.9%) |
| Perceived Stress | Asian | 23230 | 728 (3.1%) |
| Perceived Stress | More than one population | 15220 | 369 (2.4%) |
| Religious Attendance | Black or African American | 6583 | 942 (14.3%) |
| Religious Attendance | White | 67545 | 8405 (12.4%) |
| Religious Attendance | Middle Eastern or North African | 383 | 46 (12.0%) |
| Religious Attendance | Other | 9616 | 1039 (10.8%) |
| Religious Attendance | More than one population | 1522 | 114 (7.5%) |
| Religious Attendance | Asian | 2323 | 165 (7.1%) |
| Religious Spirituality | Black or African American | 39498 | 1127 (2.9%) |
| Religious Spirituality | Other | 57696 | 1502 (2.6%) |
| Religious Spirituality | Middle Eastern or North African | 2298 | 58 (2.5%) |
| Religious Spirituality | White | 405270 | 5522 (1.4%) |
| Religious Spirituality | More than one population | 9132 | 120 (1.3%) |
| Religious Spirituality | Asian | 13938 | 180 (1.3%) |
| Social Support | Black or African American | 52664 | 1987 (3.8%) |
| Social Support | Other | 76928 | 2744 (3.6%) |
| Social Support | Middle Eastern or North African | 3064 | 94 (3.1%) |
| Social Support | White | 540360 | 10638 (2.0%) |
| Social Support | Asian | 18584 | 333 (1.8%) |
| Social Support | More than one population | 12176 | 144 (1.2%) |

**Table S5:** Haseman–Elston heritability estimates across SFOH SSI adjustment models.

| Trait | Model | $h^2$ | SE | 95% CI | $p$ |
| --- | --- | --- | --- | --- | --- |
| **Anthropometric and Physiological Measures** |  |  |  |  |  |
| Waist Circumference | 1. No Adjustment | 0.339 | 0.025 | [0.289, 0.389] | 8.5e-41 |
| Waist Circumference | 2. Lead Measures | 0.345 | 0.026 | [0.294, 0.397] | 1.2e-39 |
| Waist Circumference | 4. No Adjustment + 7 Genetic PCs | 0.033 | 0.004 | [0.025, 0.040] | 8.4e-17 |
| Waist Circumference | 5. Lead Measures + 7 Genetic PCs | 0.031 | 0.004 | [0.023, 0.039] | 1.2e-14 |
| Hip Circumference | 1. No Adjustment | 0.302 | 0.024 | [0.256, 0.349] | 7.1e-37 |
| Hip Circumference | 2. Lead Measures | 0.308 | 0.025 | [0.260, 0.357] | 1.3e-35 |
| Hip Circumference | 4. No Adjustment + 7 Genetic PCs | 0.031 | 0.004 | [0.023, 0.038] | 4.8e-15 |
| Hip Circumference | 5. Lead Measures + 7 Genetic PCs | 0.029 | 0.004 | [0.022, 0.037] | 1.5e-13 |
| Body Mass Index | 1. No Adjustment | 0.268 | 0.020 | [0.228, 0.308] | 3.7e-39 |
| Body Mass Index | 2. Lead Measures | 0.269 | 0.021 | [0.228, 0.310] | 2.8e-38 |
| Body Mass Index | 4. No Adjustment + 7 Genetic PCs | 0.036 | 0.004 | [0.029, 0.043] | 2.0e-23 |
| Body Mass Index | 5. Lead Measures + 7 Genetic PCs | 0.035 | 0.004 | [0.028, 0.042] | 9.9e-22 |
| Diastolic BP | 1. No Adjustment | 0.084 | 0.012 | [0.060, 0.109] | 1.4e-11 |
| Diastolic BP | 2. Lead Measures | 0.086 | 0.013 | [0.061, 0.112] | 1.5e-11 |
| Diastolic BP | 4. No Adjustment + 7 Genetic PCs | 0.012 | 0.003 | [0.006, 0.019] | 0.00028 |
| Diastolic BP | 5. Lead Measures + 7 Genetic PCs | 0.013 | 0.003 | [0.006, 0.020] | 0.00026 |
| Heart Rate | 1. No Adjustment | 0.056 | 0.010 | [0.036, 0.076] | 2.9e-08 |
| Heart Rate | 2. Lead Measures | 0.058 | 0.011 | [0.036, 0.079] | 1.2e-07 |
| Heart Rate | 4. No Adjustment + 7 Genetic PCs | 0.012 | 0.003 | [0.005, 0.019] | 0.00038 |
| Heart Rate | 5. Lead Measures + 7 Genetic PCs | 0.011 | 0.003 | [0.005, 0.018] | 0.00098 |
| Systolic BP | 1. No Adjustment | 0.048 | 0.010 | [0.028, 0.068] | 3.8e-06 |
| Systolic BP | 2. Lead Measures | 0.047 | 0.010 | [0.027, 0.068] | 4.7e-06 |
| Systolic BP | 4. No Adjustment + 7 Genetic PCs | 0.009 | 0.003 | [0.002, 0.016] | 0.00720 |
| Systolic BP | 5. Lead Measures + 7 Genetic PCs | 0.009 | 0.003 | [0.002, 0.015] | 0.01097 |
| **Blood Chemistry and Electrolytes** |  |  |  |  |  |
| HDL Cholesterol | 1. No Adjustment | 0.274 | 0.034 | [0.208, 0.340] | 3.9e-16 |
| HDL Cholesterol | 2. Lead Measures | 0.277 | 0.034 | [0.209, 0.344] | 1.0e-15 |
| HDL Cholesterol | 4. No Adjustment + 7 Genetic PCs | 0.065 | 0.007 | [0.050, 0.079] | 5.2e-18 |
| HDL Cholesterol | 5. Lead Measures + 7 Genetic PCs | 0.065 | 0.008 | [0.050, 0.080] | 1.6e-17 |
| Glucose | 1. No Adjustment | 0.122 | 0.020 | [0.082, 0.161] | 1.9e-09 |
| Glucose | 2. Lead Measures | 0.120 | 0.021 | [0.080, 0.161] | 4.7e-09 |
| Glucose | 4. No Adjustment + 7 Genetic PCs | 0.013 | 0.006 | [0.002, 0.023] | 0.02503 |
| Glucose | 5. Lead Measures + 7 Genetic PCs | 0.011 | 0.006 | [0.000, 0.023] | 0.04344 |
| Carbon Dioxide | 1. No Adjustment | 0.103 | 0.021 | [0.062, 0.144] | 6.9e-07 |
| Carbon Dioxide | 2. Lead Measures | 0.102 | 0.021 | [0.061, 0.143] | 1.1e-06 |
| Carbon Dioxide | 4. No Adjustment + 7 Genetic PCs | 0.004 | 0.007 | [-0.009, 0.017] | 0.52463 |
| Carbon Dioxide | 5. Lead Measures + 7 Genetic PCs | 0.004 | 0.007 | [-0.009, 0.017] | 0.52583 |
| Sodium | 1. No Adjustment | 0.085 | 0.018 | [0.051, 0.120] | 1.2e-06 |
| Sodium | 2. Lead Measures | 0.086 | 0.018 | [0.051, 0.122] | 1.7e-06 |
| Sodium | 4. No Adjustment + 7 Genetic PCs | 0.008 | 0.005 | [-0.002, 0.019] | 0.13258 |
| Sodium | 5. Lead Measures + 7 Genetic PCs | 0.007 | 0.006 | [-0.003, 0.018] | 0.17746 |
| Protein | 1. No Adjustment | 0.058 | 0.015 | [0.029, 0.086] | 7.4e-05 |
| Protein | 2. Lead Measures | 0.056 | 0.014 | [0.029, 0.084] | 6.2e-05 |
| Protein | 4. No Adjustment + 7 Genetic PCs | 0.023 | 0.006 | [0.011, 0.034] | 0.00011 |
| Protein | 5. Lead Measures + 7 Genetic PCs | 0.023 | 0.006 | [0.011, 0.035] | 0.00011 |
| Chloride | 1. No Adjustment | 0.054 | 0.012 | [0.030, 0.078] | 1.2e-05 |
| Chloride | 2. Lead Measures | 0.054 | 0.012 | [0.029, 0.078] | 1.5e-05 |
| Chloride | 4. No Adjustment + 7 Genetic PCs | 0.013 | 0.006 | [0.002, 0.024] | 0.01724 |
| Chloride | 5. Lead Measures + 7 Genetic PCs | 0.014 | 0.006 | [0.002, 0.025] | 0.01674 |
| **Organ Function and Blood Cell Markers** |  |  |  |  |  |
| Urea Nitrogen | 1. No Adjustment | 0.313 | 0.040 | [0.235, 0.391] | 2.5e-15 |
| Urea Nitrogen | 2. Lead Measures | 0.316 | 0.040 | [0.237, 0.394] | 3.4e-15 |
| Urea Nitrogen | 4. No Adjustment + 7 Genetic PCs | -0.004 | 0.006 | [-0.015, 0.007] | 0.44278 |
| Urea Nitrogen | 5. Lead Measures + 7 Genetic PCs | -0.004 | 0.006 | [-0.015, 0.007] | 0.43995 |
| Alkaline Phosphatase | 1. No Adjustment | 0.097 | 0.017 | [0.063, 0.131] | 2.3e-08 |
| Alkaline Phosphatase | 2. Lead Measures | 0.093 | 0.017 | [0.060, 0.127] | 4.8e-08 |
| Alkaline Phosphatase | 4. No Adjustment + 7 Genetic PCs | 0.018 | 0.006 | [0.007, 0.030] | 0.00190 |
| Alkaline Phosphatase | 5. Lead Measures + 7 Genetic PCs | 0.017 | 0.006 | [0.005, 0.029] | 0.00568 |
| Creatinine | 1. No Adjustment | 0.075 | 0.017 | [0.043, 0.108] | 6.7e-06 |
| Creatinine | 2. Lead Measures | 0.077 | 0.017 | [0.044, 0.109] | 4.9e-06 |
| Creatinine | 4. No Adjustment + 7 Genetic PCs | 0.014 | 0.006 | [0.003, 0.025] | 0.01429 |

Table S5 – continued from previous page
| **Trait** | **Model** | **$h^2$** | **SE** | **95% CI** | **$p$** |
| --- | --- | --- | --- | --- | --- |
| Creatinine | 5. Lead Measures + 7 Genetic PCs | 0.014 | 0.006 | [0.003, 0.025] | 0.01318 |
| Alanine Aminotransferase | 1. No Adjustment | 0.072 | 0.018 | [0.037, 0.107] | 4.7e-05 |
| Alanine Aminotransferase | 2. Lead Measures | 0.071 | 0.018 | [0.036, 0.106] | 5.9e-05 |
| Alanine Aminotransferase | 4. No Adjustment + 7 Genetic PCs | 0.004 | 0.006 | [-0.008, 0.017] | 0.48976 |
| Alanine Aminotransferase | 5. Lead Measures + 7 Genetic PCs | 0.005 | 0.007 | [-0.008, 0.018] | 0.45400 |
| Aspartate Aminotransferase | 1. No Adjustment | 0.022 | 0.011 | [0.001, 0.043] | 0.04466 |
| Aspartate Aminotransferase | 2. Lead Measures | 0.022 | 0.011 | [0.001, 0.043] | 0.04033 |
| Aspartate Aminotransferase | 4. No Adjustment + 7 Genetic PCs | -0.005 | 0.006 | [-0.017, 0.007] | 0.39557 |
| Aspartate Aminotransferase | 5. Lead Measures + 7 Genetic PCs | -0.004 | 0.006 | [-0.016, 0.008] | 0.47921 |
| Hematocrit | 1. No Adjustment | 0.020 | 0.011 | [-0.001, 0.041] | 0.06152 |
| Hematocrit | 2. Lead Measures | 0.020 | 0.011 | [-0.002, 0.041] | 0.07007 |
| Hematocrit | 4. No Adjustment + 7 Genetic PCs | 0.004 | 0.006 | [-0.008, 0.016] | 0.53741 |
| Hematocrit | 5. Lead Measures + 7 Genetic PCs | 0.003 | 0.006 | [-0.009, 0.016] | 0.61479 |

**Table S6:** Haseman–Elston heritability estimates across models with SFOH MCA adjustment.

| Trait | Model | $h^2$ | SE | 95% CI | $p$ |
| --- | --- | --- | --- | --- | --- |
| **Anthropometric and Physiological Measures** |  |  |  |  |  |
| Waist Circumference | 1. No Adjustment | 0.339 | 0.025 | [0.289, 0.389] | 8.5e-41 |
| Waist Circumference | 2. Lead Measures | 0.232 | 0.021 | [0.192, 0.272] | 2.5e-29 |
| Waist Circumference | 3. Comprehensive Measures | 0.188 | 0.018 | [0.152, 0.224] | 1.4e-24 |
| Waist Circumference | 4. No Adjustment + 7 Genetic PCs | 0.033 | 0.004 | [0.025, 0.040] | 8.4e-17 |
| Waist Circumference | 5. Lead Measures + 7 Genetic PCs | 0.029 | 0.004 | [0.021, 0.037] | 9.1e-14 |
| Waist Circumference | 6. Comprehensive Measures + 7 Genetic PCs | 0.028 | 0.004 | [0.020, 0.035] | 1.4e-12 |
| Hip Circumference | 1. No Adjustment | 0.302 | 0.024 | [0.256, 0.349] | 7.1e-37 |
| Hip Circumference | 2. Lead Measures | 0.209 | 0.020 | [0.170, 0.247] | 1.7e-26 |
| Hip Circumference | 3. Comprehensive Measures | 0.165 | 0.017 | [0.132, 0.199] | 3.7e-22 |
| Hip Circumference | 4. No Adjustment + 7 Genetic PCs | 0.031 | 0.004 | [0.023, 0.038] | 4.8e-15 |
| Hip Circumference | 5. Lead Measures + 7 Genetic PCs | 0.028 | 0.004 | [0.020, 0.036] | 7.8e-13 |
| Hip Circumference | 6. Comprehensive Measures + 7 Genetic PCs | 0.027 | 0.004 | [0.019, 0.034] | 5.2e-12 |
| Body Mass Index | 1. No Adjustment | 0.268 | 0.020 | [0.228, 0.308] | 3.7e-39 |
| Body Mass Index | 2. Lead Measures | 0.155 | 0.015 | [0.126, 0.185] | 2.3e-25 |
| Body Mass Index | 3. Comprehensive Measures | 0.112 | 0.012 | [0.088, 0.136] | 1.7e-19 |
| Body Mass Index | 4. No Adjustment + 7 Genetic PCs | 0.036 | 0.004 | [0.029, 0.043] | 2.0e-23 |
| Body Mass Index | 5. Lead Measures + 7 Genetic PCs | 0.032 | 0.004 | [0.025, 0.039] | 5.5e-19 |
| Body Mass Index | 6. Comprehensive Measures + 7 Genetic PCs | 0.029 | 0.004 | [0.022, 0.036] | 1.5e-16 |
| Diastolic BP | 1. No Adjustment | 0.084 | 0.012 | [0.060, 0.109] | 1.4e-11 |
| Diastolic BP | 2. Lead Measures | 0.066 | 0.011 | [0.044, 0.087] | 1.5e-09 |
| Diastolic BP | 3. Comprehensive Measures | 0.057 | 0.010 | [0.038, 0.077] | 1.1e-08 |
| Diastolic BP | 4. No Adjustment + 7 Genetic PCs | 0.012 | 0.003 | [0.006, 0.019] | 0.00028 |
| Diastolic BP | 5. Lead Measures + 7 Genetic PCs | 0.012 | 0.003 | [0.005, 0.018] | 0.00056 |
| Diastolic BP | 6. Comprehensive Measures + 7 Genetic PCs | 0.011 | 0.003 | [0.005, 0.018] | 0.00081 |
| Systolic BP | 1. No Adjustment | 0.048 | 0.010 | [0.028, 0.068] | 3.8e-06 |
| Systolic BP | 2. Lead Measures | 0.037 | 0.009 | [0.019, 0.055] | 4.3e-05 |
| Systolic BP | 3. Comprehensive Measures | 0.031 | 0.008 | [0.014, 0.047] | 0.00021 |
| Systolic BP | 4. No Adjustment + 7 Genetic PCs | 0.009 | 0.003 | [0.002, 0.016] | 0.00720 |
| Systolic BP | 5. Lead Measures + 7 Genetic PCs | 0.009 | 0.003 | [0.002, 0.015] | 0.00901 |
| Systolic BP | 6. Comprehensive Measures + 7 Genetic PCs | 0.008 | 0.003 | [0.002, 0.015] | 0.01306 |
| Heart Rate | 1. No Adjustment | 0.056 | 0.010 | [0.036, 0.076] | 2.9e-08 |
| Heart Rate | 2. Lead Measures | 0.051 | 0.010 | [0.031, 0.071] | 3.5e-07 |
| Heart Rate | 3. Comprehensive Measures | 0.043 | 0.009 | [0.025, 0.061] | 1.6e-06 |
| Heart Rate | 4. No Adjustment + 7 Genetic PCs | 0.012 | 0.003 | [0.005, 0.019] | 0.00038 |
| Heart Rate | 5. Lead Measures + 7 Genetic PCs | 0.011 | 0.003 | [0.005, 0.018] | 0.00097 |
| Heart Rate | 6. Comprehensive Measures + 7 Genetic PCs | 0.011 | 0.003 | [0.004, 0.018] | 0.00107 |
| **Blood Chemistry and Electrolytes** |  |  |  |  |  |
| HDL Cholesterol | 1. No Adjustment | 0.274 | 0.034 | [0.208, 0.340] | 3.9e-16 |
| HDL Cholesterol | 2. Lead Measures | 0.178 | 0.025 | [0.128, 0.228] | 3.1e-12 |
| HDL Cholesterol | 3. Comprehensive Measures | 0.135 | 0.021 | [0.094, 0.176] | 8.8e-11 |
| HDL Cholesterol | 4. No Adjustment + 7 Genetic PCs | 0.065 | 0.007 | [0.050, 0.079] | 5.2e-18 |
| HDL Cholesterol | 5. Lead Measures + 7 Genetic PCs | 0.061 | 0.007 | [0.047, 0.076] | 2.2e-16 |
| HDL Cholesterol | 6. Comprehensive Measures + 7 Genetic PCs | 0.059 | 0.007 | [0.045, 0.074] | 1.5e-15 |
| Glucose | 1. No Adjustment | 0.122 | 0.020 | [0.082, 0.161] | 1.9e-09 |
| Glucose | 2. Lead Measures | 0.081 | 0.016 | [0.049, 0.113] | 8.7e-07 |
| Glucose | 3. Comprehensive Measures | 0.069 | 0.015 | [0.039, 0.099] | 5.6e-06 |
| Glucose | 4. No Adjustment + 7 Genetic PCs | 0.013 | 0.006 | [0.002, 0.023] | 0.02503 |
| Glucose | 5. Lead Measures + 7 Genetic PCs | 0.010 | 0.006 | [-0.000, 0.021] | 0.06119 |
| Glucose | 6. Comprehensive Measures + 7 Genetic PCs | 0.009 | 0.006 | [-0.001, 0.020] | 0.08829 |
| Carbon Dioxide | 1. No Adjustment | 0.103 | 0.021 | [0.062, 0.144] | 6.9e-07 |
| Carbon Dioxide | 2. Lead Measures | 0.100 | 0.021 | [0.060, 0.141] | 1.0e-06 |
| Carbon Dioxide | 3. Comprehensive Measures | 0.094 | 0.020 | [0.055, 0.133] | 2.4e-06 |
| Carbon Dioxide | 4. No Adjustment + 7 Genetic PCs | 0.004 | 0.007 | [-0.009, 0.017] | 0.52463 |
| Carbon Dioxide | 5. Lead Measures + 7 Genetic PCs | 0.004 | 0.007 | [-0.009, 0.017] | 0.54691 |
| Carbon Dioxide | 6. Comprehensive Measures + 7 Genetic PCs | 0.003 | 0.007 | [-0.009, 0.016] | 0.60079 |
| Sodium | 1. No Adjustment | 0.085 | 0.018 | [0.051, 0.120] | 1.2e-06 |
| Sodium | 2. Lead Measures | 0.068 | 0.016 | [0.037, 0.098] | 1.3e-05 |
| Sodium | 3. Comprehensive Measures | 0.053 | 0.014 | [0.026, 0.080] | 0.00011 |
| Sodium | 4. No Adjustment + 7 Genetic PCs | 0.008 | 0.005 | [-0.002, 0.019] | 0.13258 |
| Sodium | 5. Lead Measures + 7 Genetic PCs | 0.008 | 0.005 | [-0.003, 0.019] | 0.14003 |

Table S6 – continued from previous page
| Trait | Model | $h^2$ | SE | 95% CI | $p$ |
| --- | --- | --- | --- | --- | --- |
| Sodium | 6. Comprehensive Measures + 7 Genetic PCs | 0.007 | 0.005 | [-0.003, 0.018] | 0.17269 |
| Protein | 1. No Adjustment | 0.058 | 0.015 | [0.029, 0.086] | 7.4e-05 |
| Protein | 2. Lead Measures | 0.057 | 0.014 | [0.029, 0.085] | 6.6e-05 |
| Protein | 3. Comprehensive Measures | 0.056 | 0.014 | [0.028, 0.084] | 7.5e-05 |
| Protein | 4. No Adjustment + 7 Genetic PCs | 0.023 | 0.006 | [0.011, 0.034] | 0.00011 |
| Protein | 5. Lead Measures + 7 Genetic PCs | 0.023 | 0.006 | [0.011, 0.034] | 0.00012 |
| Protein | 6. Comprehensive Measures + 7 Genetic PCs | 0.023 | 0.006 | [0.011, 0.034] | 0.00012 |
| Chloride | 1. No Adjustment | 0.054 | 0.012 | [0.030, 0.078] | 1.2e-05 |
| Chloride | 2. Lead Measures | 0.054 | 0.012 | [0.030, 0.078] | 1.2e-05 |
| Chloride | 3. Comprehensive Measures | 0.052 | 0.012 | [0.028, 0.076] | 1.6e-05 |
| Chloride | 4. No Adjustment + 7 Genetic PCs | 0.013 | 0.006 | [0.002, 0.024] | 0.01724 |
| Chloride | 5. Lead Measures + 7 Genetic PCs | 0.014 | 0.006 | [0.003, 0.025] | 0.01540 |
| Chloride | 6. Comprehensive Measures + 7 Genetic PCs | 0.013 | 0.006 | [0.002, 0.024] | 0.01787 |
| **Organ Function and Blood Cell Markers** |  |  |  |  |  |
| Urea Nitrogen | 1. No Adjustment | 0.313 | 0.040 | [0.235, 0.391] | 2.5e-15 |
| Urea Nitrogen | 2. Lead Measures | 0.319 | 0.040 | [0.240, 0.397] | 1.4e-15 |
| Urea Nitrogen | 3. Comprehensive Measures | 0.324 | 0.040 | [0.245, 0.403] | 8.0e-16 |
| Urea Nitrogen | 4. No Adjustment + 7 Genetic PCs | -0.004 | 0.006 | [-0.015, 0.007] | 0.44278 |
| Urea Nitrogen | 5. Lead Measures + 7 Genetic PCs | -0.004 | 0.006 | [-0.015, 0.006] | 0.42682 |
| Urea Nitrogen | 6. Comprehensive Measures + 7 Genetic PCs | -0.005 | 0.006 | [-0.015, 0.006] | 0.39627 |
| Alkaline Phosphatase | 1. No Adjustment | 0.097 | 0.017 | [0.063, 0.131] | 2.3e-08 |
| Alkaline Phosphatase | 2. Lead Measures | 0.074 | 0.014 | [0.046, 0.103] | 2.6e-07 |
| Alkaline Phosphatase | 3. Comprehensive Measures | 0.068 | 0.014 | [0.041, 0.095] | 7.0e-07 |
| Alkaline Phosphatase | 4. No Adjustment + 7 Genetic PCs | 0.018 | 0.006 | [0.007, 0.030] | 0.00190 |
| Alkaline Phosphatase | 5. Lead Measures + 7 Genetic PCs | 0.017 | 0.006 | [0.006, 0.029] | 0.00339 |
| Alkaline Phosphatase | 6. Comprehensive Measures + 7 Genetic PCs | 0.017 | 0.006 | [0.006, 0.029] | 0.00374 |
| Creatinine | 1. No Adjustment | 0.075 | 0.017 | [0.043, 0.108] | 6.7e-06 |
| Creatinine | 2. Lead Measures | 0.077 | 0.017 | [0.044, 0.110] | 4.9e-06 |
| Creatinine | 3. Comprehensive Measures | 0.087 | 0.018 | [0.051, 0.123] | 1.7e-06 |
| Creatinine | 4. No Adjustment + 7 Genetic PCs | 0.014 | 0.006 | [0.003, 0.025] | 0.01429 |
| Creatinine | 5. Lead Measures + 7 Genetic PCs | 0.013 | 0.006 | [0.002, 0.024] | 0.01646 |
| Creatinine | 6. Comprehensive Measures + 7 Genetic PCs | 0.013 | 0.006 | [0.002, 0.024] | 0.02050 |
| Alanine Aminotransferase | 1. No Adjustment | 0.072 | 0.018 | [0.037, 0.107] | 4.7e-05 |
| Alanine Aminotransferase | 2. Lead Measures | 0.054 | 0.016 | [0.024, 0.085] | 0.00047 |
| Alanine Aminotransferase | 3. Comprehensive Measures | 0.044 | 0.014 | [0.016, 0.072] | 0.00181 |
| Alanine Aminotransferase | 4. No Adjustment + 7 Genetic PCs | 0.004 | 0.006 | [-0.008, 0.017] | 0.48976 |
| Alanine Aminotransferase | 5. Lead Measures + 7 Genetic PCs | 0.004 | 0.006 | [-0.009, 0.017] | 0.52661 |
| Alanine Aminotransferase | 6. Comprehensive Measures + 7 Genetic PCs | 0.004 | 0.006 | [-0.009, 0.016] | 0.56597 |
| Aspartate Aminotransferase | 1. No Adjustment | 0.022 | 0.011 | [0.001, 0.043] | 0.04466 |
| Aspartate Aminotransferase | 2. Lead Measures | 0.025 | 0.011 | [0.003, 0.048] | 0.02792 |
| Aspartate Aminotransferase | 3. Comprehensive Measures | 0.029 | 0.012 | [0.006, 0.053] | 0.01562 |
| Aspartate Aminotransferase | 4. No Adjustment + 7 Genetic PCs | -0.005 | 0.006 | [-0.017, 0.007] | 0.39557 |
| Aspartate Aminotransferase | 5. Lead Measures + 7 Genetic PCs | -0.005 | 0.006 | [-0.017, 0.007] | 0.39930 |
| Aspartate Aminotransferase | 6. Comprehensive Measures + 7 Genetic PCs | -0.005 | 0.006 | [-0.017, 0.007] | 0.39419 |
| Hematocrit | 1. No Adjustment | 0.020 | 0.011 | [-0.001, 0.041] | 0.06152 |
| Hematocrit | 2. Lead Measures | 0.022 | 0.011 | [0.000, 0.044] | 0.04819 |
| Hematocrit | 3. Comprehensive Measures | 0.024 | 0.012 | [0.001, 0.047] | 0.04001 |
| Hematocrit | 4. No Adjustment + 7 Genetic PCs | 0.004 | 0.006 | [-0.008, 0.016] | 0.53741 |
| Hematocrit | 5. Lead Measures + 7 Genetic PCs | 0.004 | 0.006 | [-0.009, 0.016] | 0.56299 |
| Hematocrit | 6. Comprehensive Measures + 7 Genetic PCs | 0.004 | 0.006 | [-0.009, 0.016] | 0.57168 |

**Table S7:**
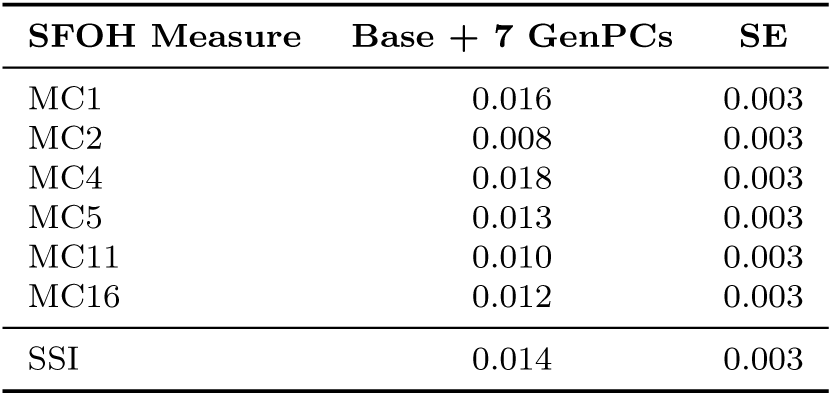
HE heritability estimates and p-values across different genetic PC adjustment models. Convergence of SFOH heritability estimates with 7 genetic PCs. Despite appearing to capture different social dimensions, all significantly heritable measures converge to *h*^2^ ≈ 0.01 − 0.02 after comprehensive genetic PC adjustment, indicating they capture the same underlying population structure rather than distinct genetic influences on different social factors.

**Table S8:**
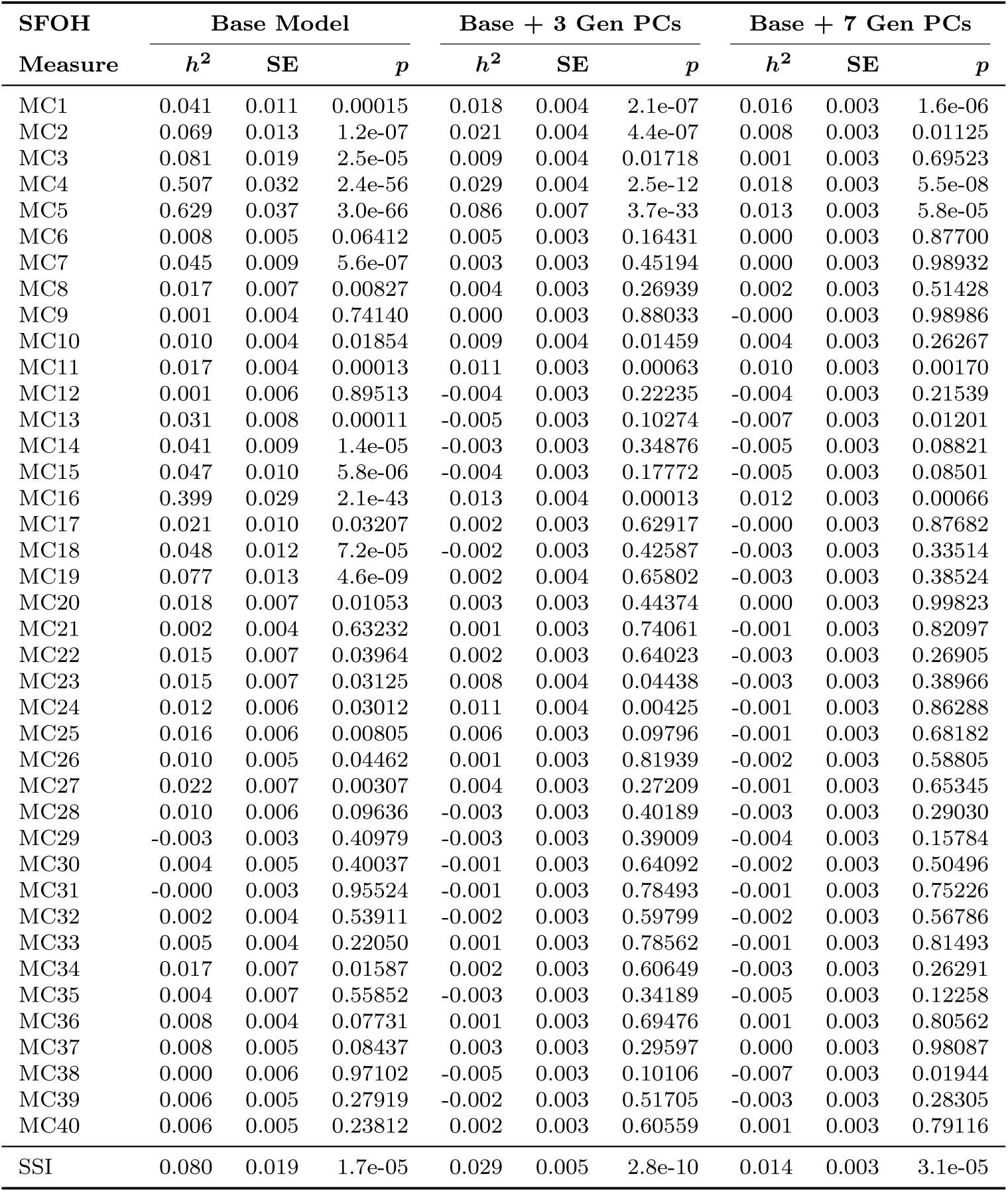
Progressive reduction in SFOH measure HE heritability with genetic PC adjustment. Heritability estimates (with respective SEs and p-values) show dramatic reductions and convergence with comprehensive PC adjustment, indicating these measures primarily capture population stratification rather than independent genetic architecture.

**Table S9:** Combined analysis group: Results of regressing SFOH MC axes onto genetic PCs. A linear combination of SFOH MCs explains non-trivial variation in genetic PC1, but limited influence on genetic PCs 2:10.

| GENPC | $R^2$ | F-stat | F p-value |
| --- | --- | --- | --- |
| GENPC1 | 0.206 | 1139.612 | 0.000 |
| GENPC2 | 0.039 | 180.670 | 0.000 |
| GENPC3 | 0.043 | 200.038 | 0.000 |
| GENPC4 | 0.031 | 141.184 | 0.000 |
| GENPC5 | 0.011 | 48.644 | 0.000 |
| GENPC6 | 0.017 | 77.088 | 0.000 |
| GENPC7 | 0.004 | 18.116 | 0.000 |
| GENPC8 | 0.008 | 36.474 | 0.000 |
| GENPC9 | 0.004 | 18.815 | 0.000 |
| GENPC10 | 0.003 | 12.976 | 0.000 |

**Table S10:**
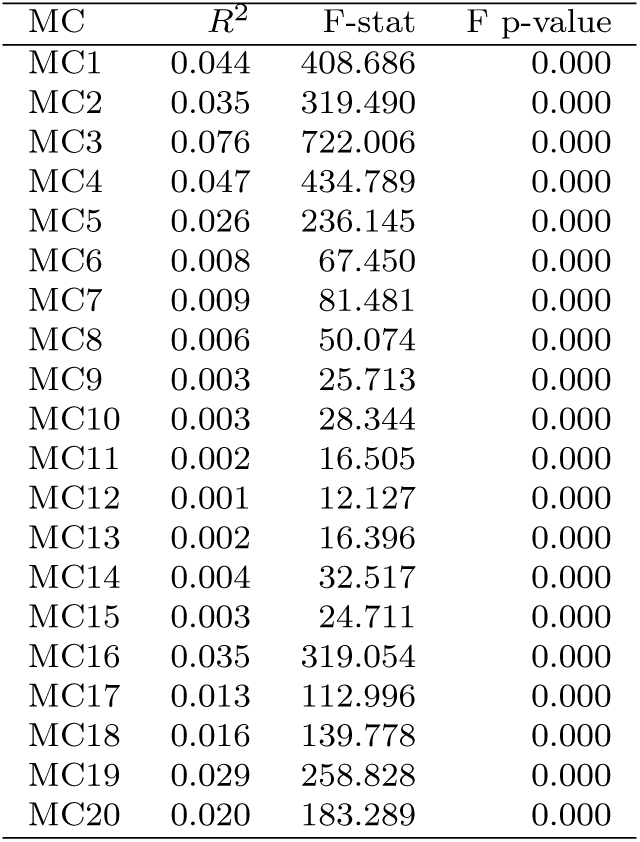
Combined analysis group: Results of regressing genetic PCs onto SFOH MC axes. Genetic PCs explain modest amounts of variation in SFOH MC axes structure with the strongest associations observed for SFOH MC3 and MC4.

## References

[1] Emil Uffelmann, Qin Qin Huang, Nchangwi Syntia Munung, Jantina de Vries, Yukinori Okada, Alicia R. Martin, Hilary C. Martin, Tuuli Lappalainen, and Danielle Posthuma. Genome-wide association studies. Nature Reviews Methods Primers, 1(1):59, August 2021. ISSN 2662-8449. doi: 10.1038/s43586-021-00056-9. Publisher: Nature Publishing Group.

[2] Jonathan Marchini, Lon R. Cardon, Michael S. Phillips, and Peter Donnelly. The effects of human population structure on large genetic association studies. Nature Genetics, 36(5):512–517, May 2004. ISSN 1546-1718. doi: 10.1038/ng1337. Publisher: Nature Publishing Group.

[3] John Novembre, Toby Johnson, Katarzyna Bryc, Zoltán Kutalik, Adam R. Boyko, Adam Auton, Amit Indap, Karen S. King, Sven Bergmann, Matthew R. Nelson, Matthew Stephens, and Carlos D. Bustamante. Genes mirror geography within Europe. Nature, 456(7218):98–101, November 2008. ISSN 1476-4687. doi: 10.1038/nature07331. Publisher: Nature Publishing Group.

[4] Daniel John Lawson, Neil Martin Davies, Simon Haworth, Bilal Ashraf, Laurence Howe, Andrew Crawford, Gibran Hemani, George Davey Smith, and Nicholas John Timpson. Is population structure in the genetic biobank era irrelevant, a challenge, or an opportunity? Human Genetics, 139(1):23–41, January 2020. ISSN 1432-1203. doi: 10.1007/s00439-019-02014-8.

[5] Kevin H. Nguyen and Megan B. Cole. Social Risk Factors, Health Insurance Coverage, and Inequities in Access to Care. American Journal of Preventive Medicine, 68(1):145–153, January 2025. ISSN 0749-3797, 1873-2607. doi: 10.1016/j.amepre.2024.09.005. Publisher: Elsevier.

[6] Mohammad Hashim Jilani, Zulqarnain Javed, Tamer Yahya, Javier Valero-Elizondo, Safi U. Khan, Bita Kash, Ron Blankstein, Salim S. Virani, Michael J. Blaha, Prachi Dubey, Adnan A. Hyder, Farhaan S. Vahidy, Miguel Cainzos-Achirica, and Khurram Nasir. Social Determinants of Health and Cardiovascular Disease: Current State and Future Directions Towards Healthcare Equity. Current Atherosclerosis Reports, 23(9):55, July 2021. ISSN 1534-6242. doi: 10.1007/s11883-021-00949-w.

[7] Cynthia A. Gómez, Dushanka V. Kleinman, Nico Pronk, Glenda L. Wrenn Gordon, Emmeline Ochiai, Carter Blakey, Ayanna Johnson, and Karen H. Brewer. Addressing Health Equity and Social Determinants of Health Through Healthy People 2030. Journal of Public Health Management and Practice, 27 (Supplement 6):S249, December 2021. ISSN 1078-4659. doi: 10.1097/PHH. 0000000000001297.

[8] Yarden S. Fraiman and Monica H. Wojcik. The influence of social determinants of health on the genetic diagnostic odyssey: who remains undiagnosed, why, and to what effect? Pediatric Research, 89(2):295–300, January 2021. ISSN 1530-0447. doi: 10.1038/s41390-020-01151-5. Publisher: Nature Publishing Group.

[9] Jimmy Phuong, Naomi O. Riches, Charisse Madlock-Brown, Deborah Duran, Luca Calzoni, Juan C. Espinoza, Gora Datta, Ramakanth Kavuluru, Nicole G. Weiskopf, Cavin K. Ward-Caviness, and Asiyah Yu Lin. Social Determinants of Health Factors for Gene–Environment COVID-19 Research: Challenges and Opportunities. Advanced Genetics, 3(2): 2100056, 2022. ISSN 2641-6573. doi: 10.1002/ggn2.202100056. eprint:https://advanced.onlinelibrary.wiley.com/doi/pdf/10.1002/ggn2.202100056.

[10] The “All of Us” Research Program | New England Journal of Medicine.

[11] Hailan Liu and Guo-Bo Chen. A novel genomic prediction method combining randomized Haseman-Elston regression with a modified algorithm for Proven and Young for large genomic data. The Crop Journal, 10(2):550–554, April 2022. ISSN 2214-5141. doi: 10.1016/j.cj.2021.09.001.

[12] Ali Pazokitoroudi, Zhengtong Liu, Andrew Dahl, Noah Zaitlen, Saharon Rosset, and Sriram Sankararaman. A scalable and robust variance components method reveals insights into the architecture of gene-environment interactions underlying complex traits. The American Journal of Human Genetics, 111(7): 1462–1480, July 2024. ISSN 0002-9297, 1537-6605. doi: 10.1016/j.ajhg.2024.05.015. Publisher: Elsevier.

[13] Suresh K. Bhavnani, Weibin Zhang, Daniel Bao, Mukaila Raji, Veronica Ajewole, Rodney Hunter, Yong-Fang Kuo, Susanne Schmidt, Monique R. Pappadis, Elise Smith, Alex Bokov, Timothy Reistetter, Shyam Visweswaran, and Brian Downer. Subtyping Social Determinants of Health in All of Us: Network Analysis and Visualization Approach. medRxiv, page 2023.01.27.23285125, August 2023. doi: 10.1101/2023.01.27.23285125.

[14] Sonali Gupta, Vincent Lam, I. King Jordan, and Leonardo Mariño-Ramírez. A composite socioeconomic deprivation index from All of Us survey data: associations with health outcomes and disparities. medRxiv, page 2024.10.04.24314904, October 2024. doi: 10.1101/2024.10.04.24314904.

[15] Jane Brown, Abhijith Biji, Kathleen Ferar, Vikas Pejaver, Eimear E. Kenny, Bian Liu, and Samira Asgari. A Scalable Framework to Integrate Social Determinants of Health into Disease Risk Models using Biobank Survey Data, September 2025. ISSN: 3067-2007 Pages: 2025.09.29.25336903.

[16] Cole Brokamp, Andrew F. Beck, Neera K. Goyal, Patrick Ryan, James M. Greenberg, and Eric S. Hall. Material community deprivation and hospital utilization during the first year of life: an urban population–based cohort study. Annals of Epidemiology, 30:37–43, February 2019. ISSN 1047-2797. doi: 10.1016/j.annepidem.2018.11.008.

[17] Arun Durvasula and Alkes L. Price. Distinct explanations underlie gene-environment interactions in the UK Biobank. The American Journal of Human Genetics, 112(3):644–658, March 2025. ISSN 0002-9297. doi: 10.1016/j.ajhg.2025.01.014. Publisher: Elsevier.

[18] Gordon K Smyth. Adjusted Likelihood Methods for Modelling Dispersion in Generalized Linear Models. Environmetricsr, 1999.

[19] Stefania Benonisdottir and Augustine Kong. Studying the genetics of participation using footprints left on the ascertained genotypes. Nature Genetics, 55(8): 1413–1420, August 2023. ISSN 1546-1718. doi: 10.1038/s41588-023-01439-2. Publisher: Nature Publishing Group.

[20] Gianmarco Mignogna, Caitlin E. Carey, Robbee Wedow, Nikolas Baya, Mattia Cordioli, Nicola Pirastu, Rino Bellocco, Kathryn Fiuza Malerbi, Michel G. Nivard, Benjamin M. Neale, Raymond K. Walters, and Andrea Ganna. Patterns of item nonresponse behaviour to survey questionnaires are systematic and associated with genetic loci. Nature Human Behaviour, 7(8):1371–1387, August 2023. ISSN 2397-3374. doi: 10.1038/s41562-023-01632-7.

[21] Shaila Musharoff, Danny Park, Andy Dahl, Joshua Galanter, Xuanyao Liu, Scott Huntsman, Celeste Eng, Esteban G. Burchard, Julien F. Ayroles, and Noah Zaitlen. Existence and implications of population variance structure, October 2018. Pages: 439661 Section: New Results.

[22] Tesfaye B. Mersha and Tilahun Abebe. Self-reported race/ethnicity in the age of genomic research: its potential impact on understanding health disparities. Human Genomics, 9(1):1, January 2015. ISSN 1479-7364. doi: 10.1186/s40246-014-0023-x.

[23] Roseann E. Peterson, Karoline Kuchenbaecker, Raymond K. Walters, Chia-Yen Chen, Alice B. Popejoy, Sathish Periyasamy, Max Lam, Conrad Iyegbe, Rona J. Strawbridge, Leslie Brick, Caitlin E. Carey, Alicia R. Martin, Jacquelyn L. Meyers, Jinni Su, Junfang Chen, Alexis C. Edwards, Allan Kalungi, Nastassja Koen, Lerato Majara, Emanuel Schwarz, Jordan W. Smoller, Eli A. Stahl, Patrick F. Sullivan, Evangelos Vassos, Bryan Mowry, Miguel L. Prieto, Alfredo Cuellar-Barboza, Tim B. Bigdeli, Howard J. Edenberg, Hailiang Huang, and Laramie E. Duncan. Genome-wide Association Studies in Ancestrally Diverse Populations: Opportunities, Methods, Pitfalls, and Recommendations. Cell, 179(3):589–603, October 2019. ISSN 0092-8674, 1097-4172. doi: 10.1016/j.cell.2019.08.051. Publisher: Elsevier.

[24] Joshua G. Schraiber and Michael D. Edge. Heritability within groups is uninformative about differences among groups: Cases from behavioral, evolutionary, and statistical genetics. Proceedings of the National Academy of Sciences, 121(12):e2319496121, March 2024. doi: 10.1073/pnas.2319496121. Publisher: Proceedings of the National Academy of Sciences.

[25] Zhaotong Lin, Souvik Seal, and Saonli Basu. Estimating SNP heritability in presence of population substructure in biobank-scale datasets. Genetics, 220 (4):iyac015, April 2022. ISSN 1943-2631. doi: 10.1093/genetics/iyac015.

[26] Jian Yang, S. Hong Lee, Michael E. Goddard, and Peter M. Visscher. GCTA: A Tool for Genome-wide Complex Trait Analysis. American Journal of Human Genetics, 88(1):76–82, January 2011. ISSN 0002-9297. doi: 10.1016/j.ajhg.2010.11.011.

[27] J. K. Haseman and R. C. Elston. The investigation of linkage between a quantitative trait and a marker locus. Behavior Genetics, 2(1):3–19, March 1972. ISSN 1573-3297. doi: 10.1007/BF01066731.

[28] Mateus H. Gouveia, Karlijn A. C. Meeks, Victor Borda, Thiago P. Leal, Fernanda S. G. Kehdy, Reagan Mogire, Ayo P. Doumatey, Eduardo Tarazona-Santos, Rick A. Kittles, Ignacio F. Mata, Timothy D. O’Connor, Adebowale A. Adeyemo, Daniel Shriner, and Charles N. Rotimi. Subcontinental genetic variation in the All of Us Research Program: Implications for biomedical research. American Journal of Human Genetics, 112(6):1286–1301, June 2025. ISSN 1537-6605. doi: 10.1016/j.ajhg.2025.04.012.

[29] Second-generation PLINK: rising to the challenge of larger and richer datasets | GigaScience | Oxford Academic.

[30] Ani Manichaikul, Josyf C. Mychaleckyj, Stephen S. Rich, Kathy Daly, Michèle Sale, and Wei-Min Chen. Robust relationship inference in genome-wide association studies. Bioinformatics, 26(22):2867–2873, November 2010. ISSN 1367-4803. doi: 10.1093/bioinformatics/btq559.

[31] Abdel Abdellaoui, Conor V. Dolan, Karin J. H. Verweij, and Michel G. Nivard. Gene–environment correlations across geographic regions affect genome-wide association studies. Nature Genetics, 54(9):1345–1354, Sep 2022. ISSN 1546-1718. doi: 10.1038/s41588-022-01158-0.

[32] Abdel Abdellaoui, David Hugh-Jones, Loic Yengo, Kathryn E. Kemper, Michel G. Nivard, Laura Veul, Yan Holtz, Brendan P. Zietsch, Timothy M. Frayling, Naomi R. Wray, Jian Yang, Karin J. H. Verweij, and Peter M. Visscher. Genetic correlates of social stratification in great britain. Nature Human Behaviour, 3(12):1332–1342, Dec 2019. ISSN 2397-3374. doi: 10.1038/s41562-019-0757-5.

[33] Laurence J. Howe, Michel G. Nivard, Tim T. Morris, Ailin F. Hansen, Humaira Rasheed, Yoonsu Cho, Geetha Chittoor, Rafael Ahlskog, Penelope A. Lind, Teemu Palviainen, Matthijs D. van der Zee, Rosa Cheesman, Massimo Mangino, Yunzhang Wang, Shuai Li, Lucija Klaric, Scott M. Ratliff, Lawrence F. Bielak, Marianne Nygaard, Alexandros Giannelis, Emily A. Willoughby, Chandra A. Reynolds, Jared V. Balbona, Ole A. Andreassen, Helga Ask, Aris Baras, Christopher R. Bauer, Dorret I. Boomsma, Archie Campbell, Harry Campbell, Zhengming Chen, Paraskevi Christofidou, Elizabeth Corfield, Christina C. Dahm, Deepika R. Dokuru, Luke M. Evans, Eco J. C. de Geus, Sudheer Giddaluru, Scott D. Gordon, K. Paige Harden, W. David Hill, Amanda Hughes, Shona M. Kerr, Yongkang Kim, Hyeokmoon Kweon, Antti Latvala, Deborah A. Lawlor, Liming Li, Kuang Lin, Per Magnus, Patrik K. E. Magnusson, Travis T. Mallard, Pekka Martikainen, Melinda C. Mills, Pål Rasmus Njølstad, John D. Overton, Nancy L. Pedersen, David J. Porteous, Jeffrey Reid, Karri Silventoinen, Melissa C. Southey, Camilla Stoltenberg, Elliot M. Tucker-Drob, Margaret J. Wright, Philipp D. Koellinger, Daniel J. Benjamin, Patrick Turley, John K. Hewitt, Matthew C. Keller, Michael C. Stallings, James J. Lee, Kaare Christensen, Sharon L. R. Kardia, Patricia A. Peyser, Jennifer A. Smith, James F. Wilson, John L. Hopper, Sara Hagg, Tim D. Spector, Jean-Baptiste Pingault, Robert Plomin, Alexandra Havdahl, Meike Bartels, Nicholas G. Martin, Sven Oskarsson, Anne E. Justice, Iona Y. Millwood, Kristian Hveem, Øyvind Naess, Cristen J. Willer, Bjørn Olav Åsvold, Jaakko Kaprio, Sarah E. Medland, Robin G. Walters, David M. Evans, George Davey Smith, Caroline Hayward, Ben Brumpton, Gibran Hemani, Neil M. Davies, George Davey Smith, Social Science Genetic Association Consortium, and Within Family Consortium. Within-sibship genome-wide association analyses decrease bias in estimates of direct genetic effects. Nature Genetics, 54(5): 581–592, May 2022. ISSN 1546-1718. doi: 10.1038/s41588-022-01062-7.

[34] Hugues Aschard, Bjarni J. Vilhjálmsson, Amit D. Joshi, Alkes L. Price, and Peter Kraft. Adjusting for Heritable Covariates Can Bias Effect Estimates in Genome-Wide Association Studies. American Journal of Human Genetics, 96 (2):329–339, February 2015. ISSN 0002-9297. doi: 10.1016/j.ajhg.2014.12.021.

[35] The All of Us Research Program Genomics Investigators. Genomic data in the All of Us Research Program. Nature, 627(8003):340–346, March 2024. ISSN 1476-4687. doi: 10.1038/s41586-023-06957-x. Publisher: Nature Publishing Group.

